# Endothelial deletion of *p53* generates transitional endothelial cells and improves lung development during neonatal hyperoxia

**DOI:** 10.1101/2024.05.07.593014

**Authors:** Lisandra Vila Ellis, Jonathan D Bywaters, Jichao Chen

**Author notes:** corresponding author, **CONTACT:** Lisandra Vila Ellis. contributed equally to this work (Lisandra Vila Ellis, Jonathan D Bywaters).

## Abstract

Bronchopulmonary dysplasia (BPD), a prevalent and chronic lung disease affecting premature newborns, results in vascular rarefaction and alveolar simplification. Although the vasculature has been recognized as a main player in this disease, the recently found capillary heterogeneity and cellular dynamics of endothelial subpopulations in BPD remain unclear. Here, we show Cap2 cells are damaged during neonatal hyperoxic injury, leading to their replacement by Cap1 cells which, in turn, significantly decline. Single-cell RNA-seq identifies the activation of numerous p53 target genes in endothelial cells, including *Cdkn1a (p21)*. While global deletion of *p53* results in worsened vasculature, endothelial-specific deletion of *p53* reverses the vascular phenotype and improves alveolar simplification during hyperoxia. This recovery is associated with the emergence of a transitional EC state, enriched for oxidative stress response genes and growth factors. These findings implicate the p53 pathway in EC type transition during injury-repair and highlights the endothelial contributions to BPD.

## INTRODUCTION

Bronchopulmonary dysplasia (BPD) is a chronic disease that develops in premature newborns following ventilation and oxygen treatment for acute respiratory failure due to their shortened gestation (1). BPD results in disrupted lung development characterized by alveolar simplification – enlarged alveoli with reduced septation, and vascular rarefaction – reduced and dysmorphic vasculature (1). Historically, much of the focus has been on the epithelium and mesenchyme as the target cells that underlie BPD pathology (2). However, limited studies propose the vasculature as a main player in the pathogenesis of BPD, something known as the “vascular hypothesis” (1). Recent studies have explored the transcriptomic changes in mouse models of BPD using single-cell RNA-sequencing (scRNA-seq) (3–5); however, the endothelial-specific contributions remain unexplored.

Using scRNA-seq, we and others have shown that lung capillaries are made up of two transcriptionally distinct endothelial cell (EC) populations – PLVAP+ ECs (also known as Cap1 or gCap) and CAR4+ ECs (Cap2, aCap or aerocytes) (6–8). These two populations also differ in morphology and localization within the alveolar space, with Cap2 cells being web-like, larger and contacting the epithelium without intervening cells. Alveolar type 1 (AT1) cells, their ultrathin epithelial counterpart in the air-blood barrier, secrete *Vascular endothelial growth factor A (Vegfa)*, a signal required for Cap2 cell specification (6). Additionally, our work revealed that Cap2 cells are located in regions undergoing secondary septation, and aid in AT1 cell folding during alveologenesis (6, 9). This critical step of alveologenesis is compromised in BPD (1), suggesting the endothelium could be a major contributor to this disease.

In this study of the endothelial-specific response in BPD, we employ a widely-used mouse model of neonatal hyperoxia coupled with EC type-specific drivers, which allows us to investigate endothelial injury. scRNA-seq analysis reveals endothelial upregulation of p53 target genes in this model, prompting us to test the function of *p53* using both a global knockout, and an endothelial-specific deletion. Strikingly, endothelial *p53* deletion followed by hyperoxic injury leads to the emergence of a previously unknown transitional EC state, associated with improvements in both vascular and alveolar phenotypes.

## RESULTS

### Neonatal hyperoxia results in decreased Cap1 vasculature

To investigate how neonatal hyperoxia disrupts the lung vasculature, we exposed newborn mouse pups to either hyperoxia (80% O_2_) or room air (21% O_2_) continuously for 14 days, without a recovery period (Fig. S1A). This is an established model of BPD with concurrent injury and repair, where injury dominates (10, 11). We evaluated epithelial and vascular integrity and found that compared to room air, mice in hyperoxia expectedly showed reduced vasculature and an enlargement of the airspace (Fig. 1A).

**Figure 1.**
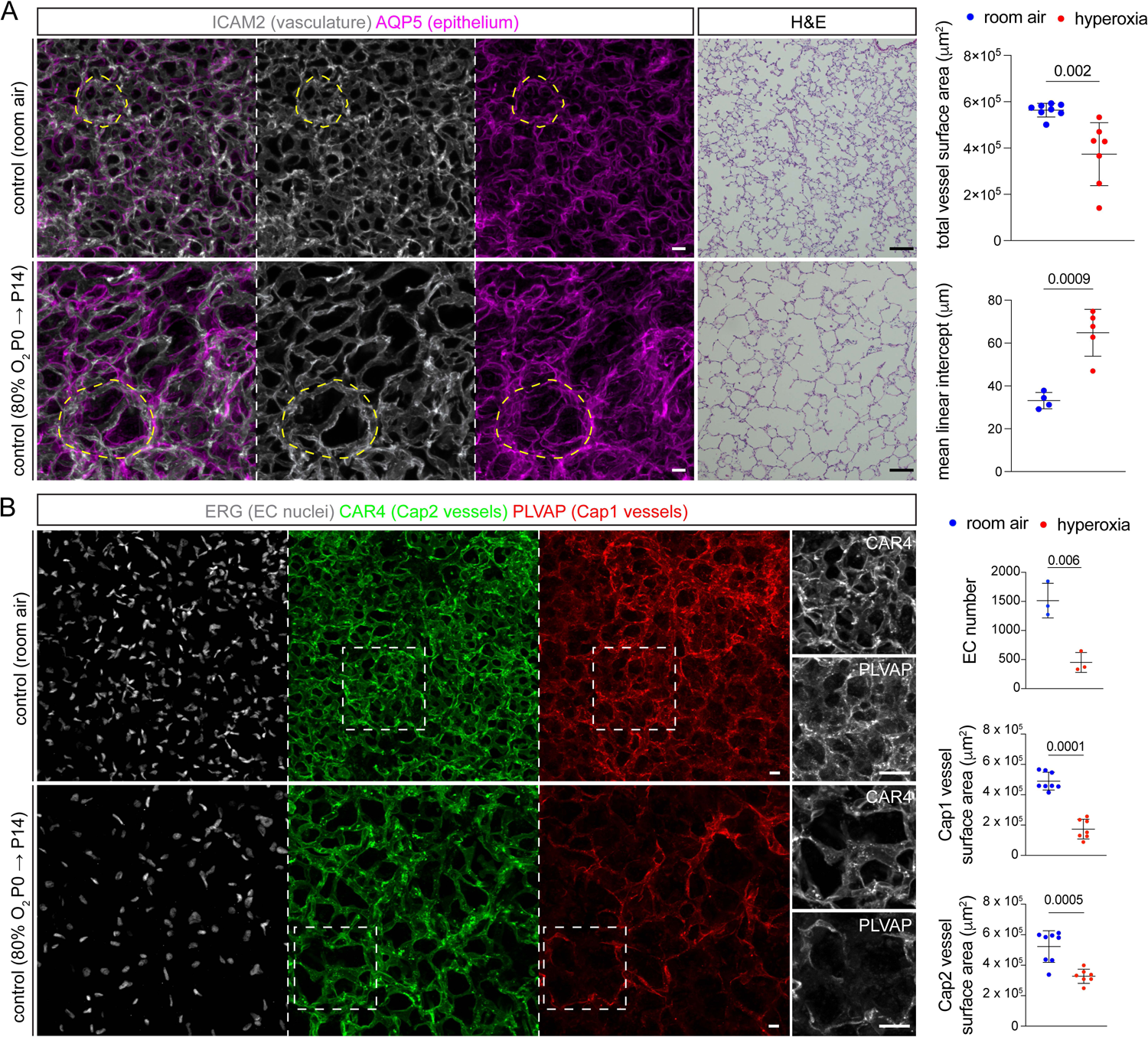
Hyperoxia exposure results in alveolar simplification with cell type-specific effects in ECs. (**A**) En-face view of immunostained lungs and H&E sections from room air and hyperoxia conditions and associated quantification. Hyperoxia-treated lungs show alveolar simplification, demonstrated by enlarged alveolar islands (dash; AQP5), and vascular rarefaction, as measured by ICAM2 staining (Student’s t test). H&E staining confirms significant enlargement of the airspace, as quantified by mean linear intercept (MLI) (Student’s t test). Each symbol represents the average of 3 distinct regions imaged within 1 mouse lung. (**B**) En-face view of immunostained lungs showing the effect of hyperoxia on distinct EC types. Boxed regions are magnified. Hyperoxia exposure results in a significant reduction in total EC number (ERG), Cap1 vasculature surface area (PLVAP), and Cap2 vasculature surface area (CAR4), with the most dramatic reduction being Cap1-specific (Student’s t test). Images are representative of at least 3 littermate pairs. P, postnatal. Scale bars, 10 um (white bars), 100 um (black bars).

To better understand the timing of Cap1/Cap2-specific injury in this model, we conducted immunostaining at various timepoints including 3, 7 and 14 days after hyperoxia (Fig. S1A). After 3 days of hyperoxia, EC number, Cap1 and Cap2 vessel area remained unchanged (Fig. S1B, C). By day 7, a reduction in EC number and Cap2 vessel area became noticeable suggesting Cap2 cells are initially affected and possibly more susceptible to hyperoxia (Fig. S1B, C). By day 14, we observed an overall reduction in EC number and total vessel area, accompanied by a preferential loss of Cap1 cell expression of its marker gene during hyperoxia, representing a decline in Cap1 vasculature (Fig. 1B). Conversely, CAR4 expression remained uniformly present in the remaining vasculature, in contrast to its patchy loss after 7 days of injury (Fig. 1B, Fig. S1B). These findings suggest EC type-specific damage and response to hyperoxia.

In patients with BPD, male newborns have shown worse outcomes compared to females (12), and these sex disparities have been explored at the transcriptional level (3, 12, 13). Therefore, we compared Cap1 and Cap2 vessel areas in male and female mice across conditions after 14 days of hyperoxia, but found no significant differences (Fig. S1D).

### Cap2 cells are damaged upon hyperoxia, and replaced by Cap1 cells, exacerbating Cap1 cell loss

To evaluate EC dynamics in this BPD model, we exposed pups pre-labeled with Cap1- or Cap2-specific drivers to hyperoxia. Using scRNA-seq, we identified *c-Kit protooncogene* (*Kit*) as a specific marker for Cap1 cells and *Car4* as a marker for Cap2 cells (Fig. S2A). We obtained a *Kit^CreERT2^* mouse and tested its recombination efficiency and specificity by crossing it to a *Rosa^Sun1GFP^* reporter (14, 15). Injection of postnatal pups with tamoxifen for 24 hours revealed it to be highly efficient (92%) and specific (100%) (Fig. S2B, C). We then coupled this driver with the reporter *Rosa^tdT^* for lineage tracing Cap1 cells (16). For Cap2 cells, lineage tracing was accomplished using our recently developed *Car4^CreER^*(17). To maximize cell recombination, a high dose of tamoxifen was administered. We implemented a 48-hour interval before hyperoxia exposure to allow sufficient time for tamoxifen clearance to (1) minimize injury-induced, non-specific driver induction and labeling and (2) mitigate heightened mortality risk during hyperoxia post-tamoxifen injection (18) (Fig. S2D). Finally, we characterized the alveolar simplification phenotype in these two models to ensure tamoxifen injection did not impact the phenotype (Fig. S2E, F) and found it to be comparable to wild type mice (Fig. 1A).

Given the reporter’s substantial accumulation in the cell nucleus, we employed tdT to quantify individual cells, simultaneously matching the nucleus with PLVAP and CAR4 antibodies as surface markers. Lineage tracing of Cap1 cells confirmed a decrease in Cap1 cell number in hyperoxia (Fig. 2A, B). Although some tdT+PLVAP+ Cap1 cells remained, we noticed that many Cap1 cells now expressed CAR4, indicating an increase in transition from Cap1 to Cap2 cells (Fig. 2A, B). We previously showed that both Cap1 and Cap2 cells arise from endothelial progenitors that closely resemble Cap1 cells (6), and this transition was corroborated by others (7). Additionally, Cap1 cells have been proposed to act as a source of new Cap2 cells upon injury and regeneration (8). While around 15% of Cap1 cells were observed transitioning to Cap2 in room air, which is expected at this early developmental stage, we observed a two-fold increase in Cap1 to Cap2 conversion in hyperoxia (Fig. 2A, B).

**Figure 2.**
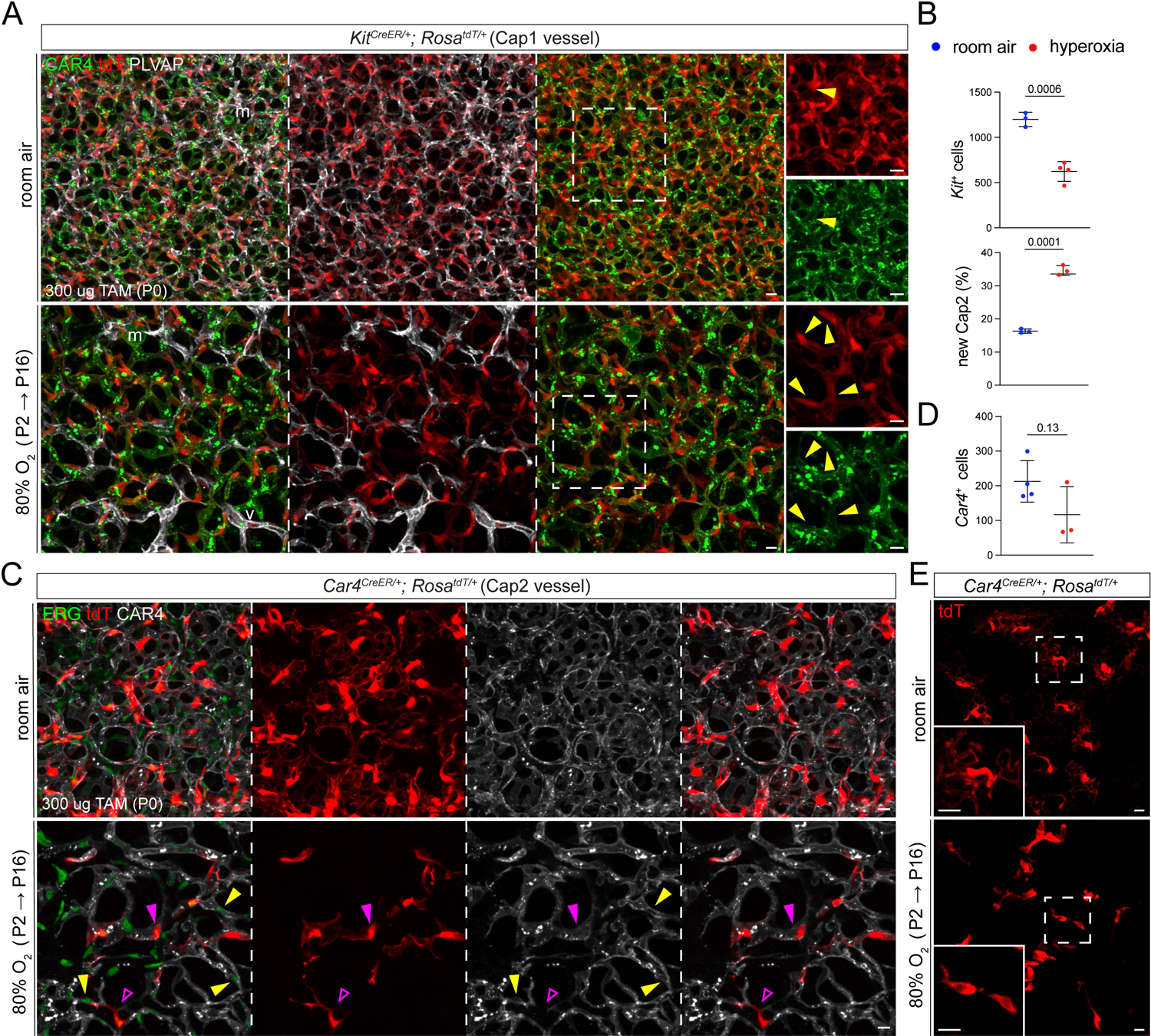
Cap1 ECs convert to Cap2 to replace cell type-specific EC loss during hyperoxia. (**A**) En-face view of immunostained lungs from lineage traced *Kit^CreER^; Rosa^tdT/+^* mice, demonstrating a significant decrease in Cap1 EC number in hyperoxia relative to room air. Hyperoxia-treated lungs also exhibited a stark increase in the proportion of tdT+ cells that express CAR4 (yellow arrowheads), suggesting increased conversion from Cap1 to Cap2 in hyperoxia. Boxed regions are magnified. (**B**) Top: quantification of tdT-positive Cap1 ECs in room air and hyperoxia, revealing a significant decrease in lineage traced Cap1 ECs (Student’s t test). Bottom: quantification of the percentage of Car4+/tdT+ cells of total tdT+ cells in room air and hyperoxia, confirming a significant increase in conversion from lineage traced Cap1 ECs to Cap2 in hyperoxia (Student’s t test). (**C**) En-face view of immunostained lungs from lineage traced *Car4^CreER^; Rosa^tdT/+^* mice. Hyperoxia exposure resulted in a reduction in tdT-positive Cap2 cells, representing Cap2 ECs present before treatment (magenta filled arrowhead). Some remaining tdT-positive cells showed a loss in CAR4 expression in hyperoxia (open arrowhead), while tdT-negative Cap2 cells (CAR4+) also appeared in hyperoxia, mostly representing new Cap2 cells converted from Cap1 (yellow arrowhead). (**D**) Quantification of tdT-positive Cap2 ECs in room air and hyperoxia, showing a downward trend in lineage traced Cap2 ECs (Student’s t test). (**E**) En-face view of immunostained lungs showing individual tdT-labeled Cap2 cells in room air and hyperoxia, demonstrating the loss of the expansive, net-like morphology of Cap2 cells upon sustained hyperoxic injury. Boxed regions are magnified. Images are representative of at least 3 littermate pairs. For quantification, each symbol represents the average of 3 distinct regions imaged within 1 mouse lung. Scale bars, 10 um. m, macrophage. v, large vessel. P, postnatal. TAM, 300 ug of tamoxifen.

This transition from Cap1 to Cap2 cells, in combination with the decrease in CAR4 expression after 7 days of hyperoxia (Fig. S1B), led us to hypothesize that Cap2 cells are susceptible to hyperoxic injury. Employing lineage tracing to label Cap2 cells before exposure to hyperoxia, we noted a substantial reduction in the number of tdT+ cells, representing original Cap2 cells (pre-injury), despite the emergence of CAR4+tdT-cells signifying new Cap2 cells (Fig. 2C, D). Additionally, some tdT+ cells in hyperoxia exhibited loss of CAR4 expression, suggesting injured Cap2 cells could lose their marker genes. Our previous work highlighted the expansive morphology of Cap2 cells, allowing them to contribute to multiple vessel segments (6). We used the same model of sparse cell labeling to visualize individual Cap2 cell morphology. We observed that, in hyperoxia, Cap2 cells deviated from their typical web-like morphology, displaying a range of shapes but all consistently smaller in size (Fig. 2E), possibly cell shrinkage prior to death.

To summarize the cellular changes, we observed that upon hyperoxia treatment (1) both Cap1 and Cap2 cells are damaged and decrease in number; (2) surviving Cap2 cells lose their distinct morphology and markers; (3) Cap1 cells transition into Cap2 cells. During normal development at room air, Cap1 cells proliferate and undergo limited conversion into Cap2 cells (6). In hyperoxia, however, Cap1 cells convert at a high rate with a net result of greater impact on Cap1 than Cap2 cells.

### Neonatal hyperoxia upregulates p53 target genes

To examine the transcriptional changes of ECs in hyperoxia, we performed scRNA-seq after 14 days of 80% O_2_ exposure and assessed a total of 2395 sorted ECs (Fig. 3A). Using established marker genes (6), each endothelial population was identified (Fig. 3B). In accordance with our previous work (6), at room air Cap1 cells represent the bulk of the lung vasculature while Cap2 cells were a small but distinct cluster (Fig. 3C, Table S1). However, in hyperoxia, scRNA-seq revealed a decrease in Cap1 cells and a proportional increase in Cap2 cells (Fig. 3C). These changes were consistent with our immunostaining and lineage tracing observations (Fig. 1B, 2A). Notably, proliferative ECs also increased in number under hyperoxia (Fig. 3C).

**Figure 3.**
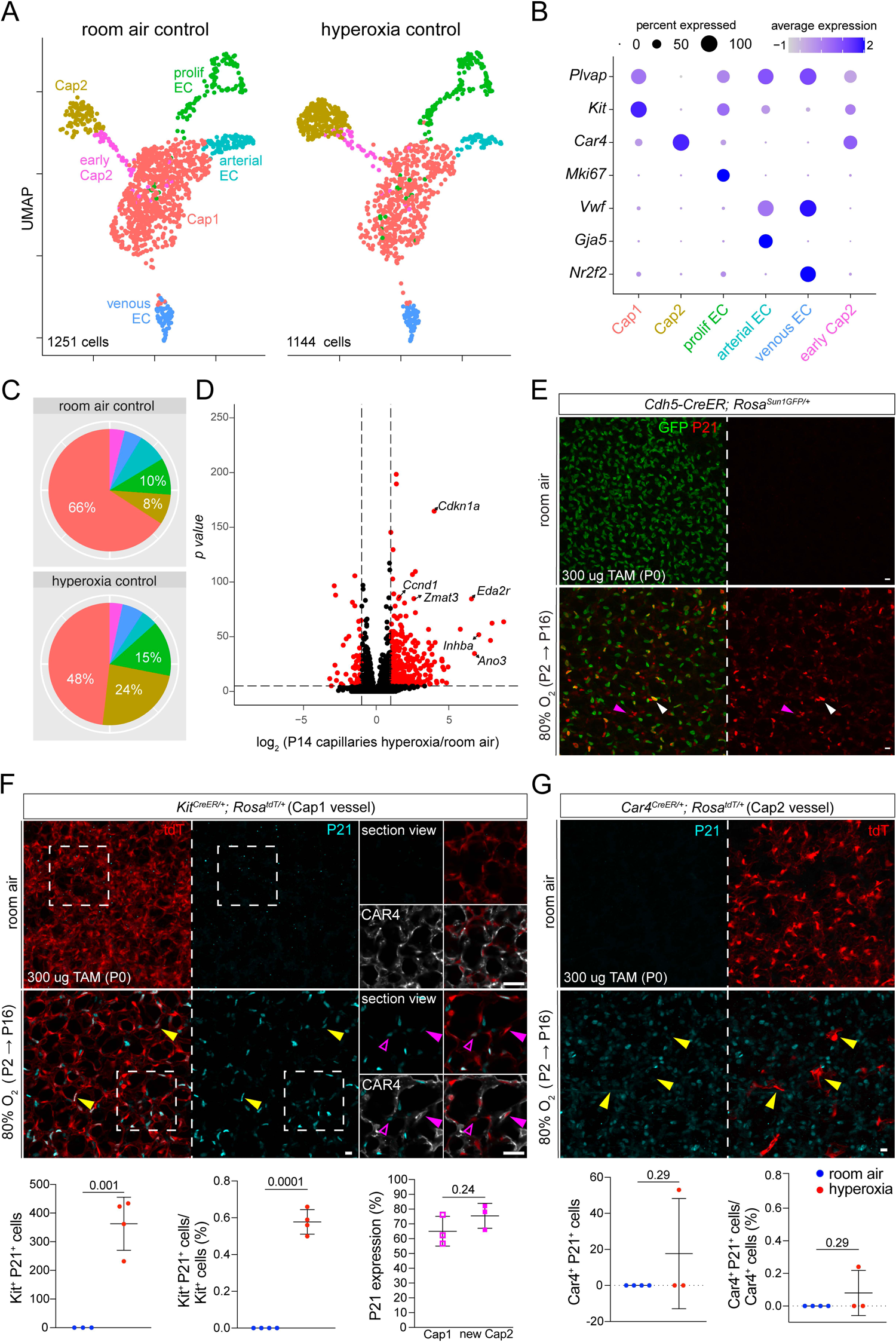
Hyperoxia exposure causes upregulation of *p53* target genes in capillary ECs. (**A**) UMAP of purified lung ECs in room air and hyperoxia. ECs are color coded according to cell population. (**B**) Dot plot showing markers used to identify each EC population. (**C**) Pie charts demonstrating the proportions of each EC population in both conditions. Hyperoxia exposure results in a substantial reduction in Cap1 cells and an increase in Cap2 cells. See Table S1 for complete analysis of cell numbers per population. (**D**) Volcano plot showing differential gene expression in capillaries between hyperoxia and room air lungs. Hyperoxia exposure causes significant upregulation of many *p53* target genes in capillary ECs. See Table S2 for the complete dataset. (**E**) En-face view of immunostained lungs showing GFP-labeled ECs in room air and hyperoxia. Hyperoxic injury causes widespread upregulation of *p53* target gene P21 in ECs (white arrowhead) and non-ECs (magenta arrowhead) compared to room air control. (**F**) En-face view of immunostained lungs showing nearly 60% of lineage traced (tdT+) Cap1 cells express P21 in hyperoxia (yellow arrowhead) and its associated quantification (below, Student’s t test). Boxed regions are magnified and shown as a section view. Both CAR4-/tdT+ Cap1 cells (representing non-converted Cap1 ECs; open magenta arrowhead) and CAR4+/tdT+ Cap2 cells (representing Cap1 cells converted into Cap2; filled magenta arrowhead) were found to express P21 with no enrichment observed in either population. (**G**) En-face view of immunostained lungs showing most lineage traced Cap2 cells were found to lack P21 expression in hyperoxia (yellow arrowhead). Images are representative of at least 3 littermate pairs. For quantification, each symbol represents the average of 3 distinct regions imaged within 1 mouse lung. P, postnatal. Scale bars, 10 um. TAM, 300 ug of tamoxifen.

Subsequently, we analyzed gene expression across the entire capillary population under both conditions and found a significant upregulation of p53 target genes, including *Cdkn1a (p21)*, a cell cycle inhibitor that protects the lung from oxidative stress (19) (Fig. 3D, S3A, Table S2). While *p53* activation in the hyperoxic lung has been documented before (20, 21), a recent publication investigating the p53/p21 pathway in hyperoxia was limited to *Von Willebrand Factor* (VWF)-positive macrovasculature, thus excluding the capillary response (22). To specifically label and trace ECs during hyperoxia, and because our EC nuclear marker (ERG) and P21 antibody were from the same species, we used a *Cdh5-CreER* mouse, a pan-endothelial driver, in combination with a *Rosa^Sun1GFP^*reporter (15). Immunofluorescence confirmed the P21 upregulation in ECs, as well as non-ECs (Fig. 3E). We also observed this upregulation by RNAscope (Fig. S3B). Further analysis of the scRNA-seq data demonstrated *Cdkn1a (p21)* upregulation in all EC types in hyperoxia compared to room air, except arteries (Fig. S3C).

To better understand capillary-specific *p53* activation, we assessed the expression of P21 in the Cap1 and Cap2 lines, *Kit^CreER^*and *Car4^CreER^*, respectively, after hyperoxia (Fig. 3F, G). Approximately 60% of Kit-positive cells (tdT+) demonstrated co-localization with P21, at a comparable frequency in Cap1 cells (tdT+CAR4-) and newly formed Cap2 cells (tdT+CAR4+) (Fig. 3F). The extent of hyperoxia-induced alveolar and vascular simplification in this model varies among mice. In samples exhibiting more severe injury as assessed by level of alveolar simplification, none of the initially labeled Cap2 cells expressed P21 (Fig. 3G). In those with milder injury, more lineage traced (tdT+) Cap2 cells persisted and few, but some of these cells expressed P21 (Fig. 3G). Therefore, Cap1 to Cap2 cell conversion is associated with *p53* activation; more of the new, Cap1-derived Cap2 cells express P21 than pre-existing Cap2 cells.

### Global deletion of *p53* worsens Cap2 vasculature in hyperoxia

To investigate the role of p53 pathway activation in hyperoxia, we obtained a *p53^flox^* mouse and crossed it with a *CMV-Cre* mouse (23, 24), inducing the deletion of the *loxP*-flanked gene in all tissues. We then selectively bred progeny lacking both the *flox* allele and the *Cre*, establishing a *p53* null mouse model. We confirmed the deletion of *p53* by immunostaining of its target gene P21 (Fig. 4A). P21 expression was absent in room air (Fig. 3E-G, Fig. 4A), whereas it was evident in the control hyperoxia group. In the *p53* null, however, P21 was not detected, indicating robust deletion of *p53* (Fig. 4A).

**Figure 4.**
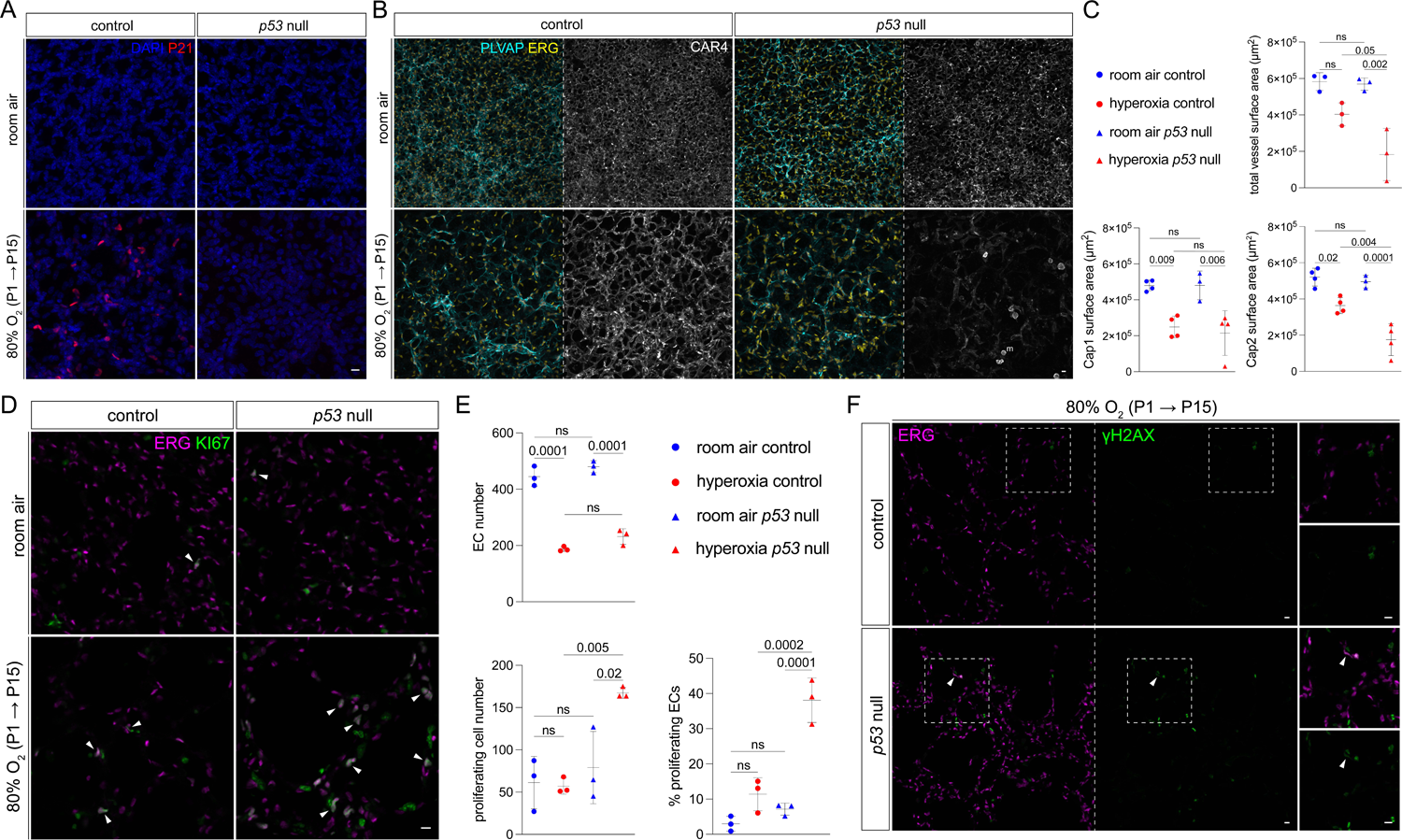
Global *p53* deletion in hyperoxia results in increased EC proliferation and Cap2-specific decrease. (**A**) En-face view of immunostained lungs demonstrating efficient deletion of *p53* in the hyperoxia *p53* null condition compared to hyperoxia control by P21 staining. (**B**) En-face view of immunostained lungs showing the effect of *p53* deletion on total EC nuclei (ERG), Cap1 vasculature (PLVAP), and Cap2 vasculature (CAR4) in room air and hyperoxia, demonstrating a decrease in CAR4 staining in the hyperoxia *p53* null. (**C**) Quantification of Cap1 surface area, Cap2 surface area, and total vessel surface area (as measured by ICAM2 staining) in each condition (1-way ANOVA with Tukey’s multiple comparisons; ns, not significant), demonstrating a significant reduction in Cap2 and total vessel area in the hyperoxia *p53* null. (**D**) Section immunostaining showing the effect of *p53* deletion and hyperoxia treatment on cell proliferation (KI67) in ECs (ERG, white arrowheads) and non-ECs. (**E**) Quantification showing hyperoxia exposure or *p53* deletion alone did not result in a significant increase in overall proliferation or EC proliferation compared to room air controls, but hyperoxia *p53* null mice exhibited significant increases in both EC-specific and overall proliferation (1-way ANOVA with Tukey’s multiple comparisons; ns, not significant). (**F**) Section immunostaining evaluating DNA damage (γH2AX) in hyperoxia-treated control and *p53* null lungs. DNA damage is higher in hyperoxia *p53* null lungs compared to hyperoxia control, with sporadic co-localization of γH2AX to ECs (ERG) in the hyperoxia *p53* null lung, indicating endothelial damage (white arrowheads). Boxed regions are magnified. Images are representative of at least 3 littermate pairs. For quantification, each symbol represents the average of 3 distinct regions imaged within 1 mouse lung. P, postnatal. m, macrophage. Scale bars, 10 um.

Assessment of the vascular phenotype, comparing control and *p53* null mice under room air vs hyperoxia conditions (Fig. 4B), revealed no differences in Cap1 and Cap2 cells between the control and *p53* null mice in room air which was expected since we did not see P21 expression in room air (Fig. 4A). In hyperoxia, the *p53* null mice exhibited a further decrease in CAR4 expression and thus Cap2 vessel area (Fig. 4C). These results could mean an increase in Cap2 cell death, damaged Cap2 cells with less marker expression, or decreased conversion from Cap1 to Cap2 fate. To further characterize the alveolar simplification phenotype normally observed in hyperoxia, we examined the alveolar surface of the lungs (Fig. S4A). Our analysis revealed that both control and *p53* null mice in hyperoxia experience similar enlargement of airspace. Quantification of the airspace by MLI did not find a significant difference between control and mutant in hyperoxia (Fig. S4A).

To dissect the cellular processes influenced by *p53* in hyperoxia within the vasculature, we examined endothelial proliferation in the *p53* null mice. In the room air control, consistent with the early developmental stage of the lungs, we observed a modest percentage of proliferative ECs, a trend that did not significantly change in hyperoxia (Fig. 4D, E). Notably, in the *p53* null mice exposed to hyperoxia, we detected a significant increase in endothelial proliferation (Fig. 4D, E). Increased proliferation could also be due to ECs escaping DNA damage checkpoints and death, unrelated to conversion, given that *p53* is known to be activated in response to DNA damage (25). We used the phosphorylated histone H2AX (γH2AX) as a DNA damage marker. In the hyperoxia control, rare non-endothelial γH2AX expression was detected, suggesting efficient cellular repair. Remarkably, *p53* null mice in hyperoxia exhibited frequent γH2AX, with rare co-localization to ECs, suggesting a potential role for *p53* activation in the regulation of DNA repair in non-ECs during hyperoxia (Fig. 4F). In addition to cell cycle, *p53* initiates apoptosis in cases of irreparable DNA damage (25).

Therefore, we evaluated cell death in hyperoxia between control and *p53* null mice (Fig. S4B). Although we observed an increase in cell death in the mutant, co-localization of γH2AX and cleaved CASPASE3 was not evident, suggesting DNA damage and repair could be sequential, where damaged cells that couldn’t undergo repair subsequently died. Taken together, this could imply that in the absence of *p53*, and in the context of hyperoxia, Cap1 cells favor a proliferation phenotype rather than a conversion/specification path, which could explain the decrease in Cap2 vessels. Additionally, the paucity of γH2AX in ECs would indicate that ECs are more resilient to DNA damage, get more quickly cleared, or are repaired faster upon hyperoxic injury.

To dissect the transcriptomic changes in the *p53* null, we conducted scRNA-seq, profiling 4471 ECs across control and mutant mice in both room air and hyperoxia (Fig. 5A). Identification of all endothelial subpopulations was achieved using previously described markers (Fig. 3B, 5B), and the proportions of each cell type were calculated (Fig. 5C, Table S1).

**Figure 5.**
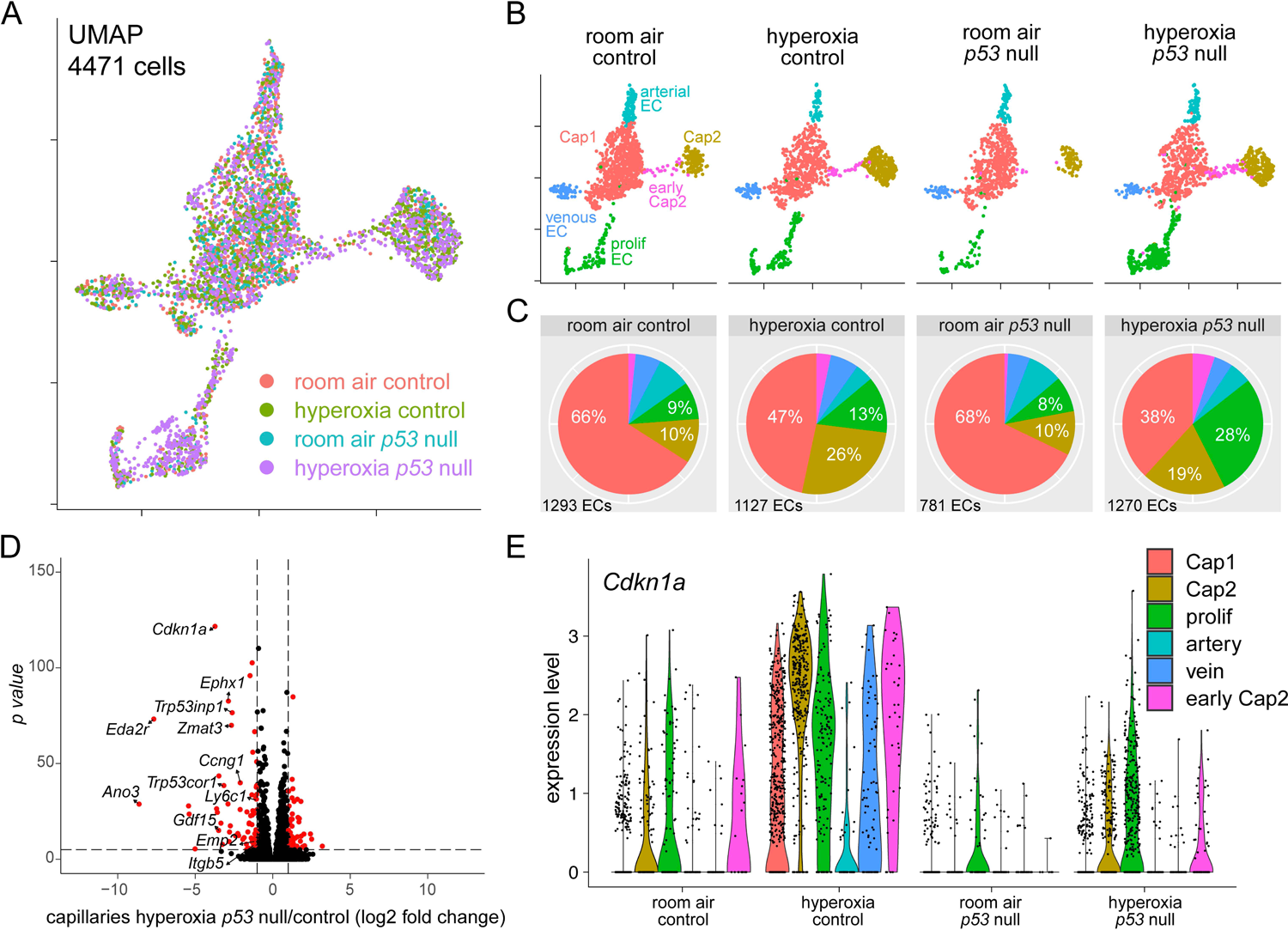
scRNA-seq reveals transcriptional changes associated to deletion of *p53* in hyperoxia-treated *p53* null lung ECs. (A) UMAP of lung ECs, color-coded and overlayed from four different experimental conditions. (B) UMAPs of lung ECs from each experimental condition, color-coded according to cell population. (**C**) Pie charts showing the proportion of each EC population in each experimental condition. Notably, proliferative ECs represent a larger proportion of ECs in the hyperoxia *p53* null lung compared to any other condition. See Table S1 for complete analysis of cell numbers per population. (**D**) Volcano plot depicting differentially expressed genes in capillaries between the hyperoxia *p53* null vs. the hyperoxia control conditions. See Table S3 for the complete dataset. (**E**) Violin plots showing expression level of p53 target *Cdkn1a* in each experimental condition, split by EC population. As compared to the hyperoxia control condition, hyperoxia *p53* null lung ECs exhibit drastically reduced *Cdkn1a* expression across all populations.

Consistent with immunostaining results, proliferative ECs substantially increased in the hyperoxia-treated *p53* null. Surprisingly, the *p53* null Cap2 cluster in hyperoxia was larger than in room air, albeit smaller than in the hyperoxia control (Fig. 5C). It is possible that Cap2 cells are present but lack expression of their mature markers such as CAR4 (Fig. 4B), making them detectable by scRNA-seq but not by staining. Additionally, smaller, more damaged or less mature Cap2 cells may be disproportionally captured during sorting. To test this, we created a list of Cap2 markers expressed in room air at P14 and used it to create a gene score to evaluate the changes in Cap2 cells across different conditions (Fig. S5A). This revealed that Cap2 cells in the *p53* null in hyperoxia downregulate their marker genes (Fig. S5B). In a closer look, we compared the top 22 Cap2 genes across Cap2 ECs in all four samples and saw a distinct downregulation in the *p53* null in hyperoxia, confirming these cells are more damaged (Fig. S5C). Differential gene analysis comparing control and *p53* null in hyperoxia revealed the expected changes associated with the deletion of *p53*, with a decline in p53 target genes such as *p21* (*Cdkn1a*), as well as some Cap2 marker genes such as *Emp2* and *Itgb5* (Fig. 5D, E, Table S3). Altogether, this shows that *p53* is directly, or indirectly through non-ECs, protective for ECs in hyperoxia.

### Conditional deletion of *p53* in the endothelium improves vascular and alveolar simplification in hyperoxia

To explore the endothelial specific role of *p53*, we conditionally deleted it using the pan-endothelial driver *Cdh5-CreER* (26). Despite efficient *p53* deletion, as indicated by P21 staining, Cap1 cells exhibited an overall improvement (Fig. S6A). However, due to variability in recombination efficiency and increased mortality associated with the combination of tamoxifen injections and hyperoxia, we opted for the use of a *Tek-Cre* (or *Tie2-Cre*) mouse instead (27), hereafter referred to as *p53^ΔEC^*. While *Tek* is enriched postnatally in Cap1 cells, it is expressed embryonically in the endothelial progenitors that give rise to both Cap1 and Cap2 (Fig. S6B); thus, this model is expected to achieve pan-endothelial deletion. Additionally, lineage tracing with the *Rosa^tdT^* reporter mouse showed that we can target both Cap1 and Cap2 cells with this driver with few escapers (Fig. S6C). Once again, since the antibodies were the same species, we employed a *Rosa^Sun1GFP^* reporter to validate the deletion by co-localizing GFP and P21 staining. We observed that most recombined cells exhibited a loss of P21 expression, except for rare escapers, which typically co-expressed CAR4 (Fig. 6A), and we also confirmed these findings via RNAscope (Fig. S6D).

**Figure 6.**
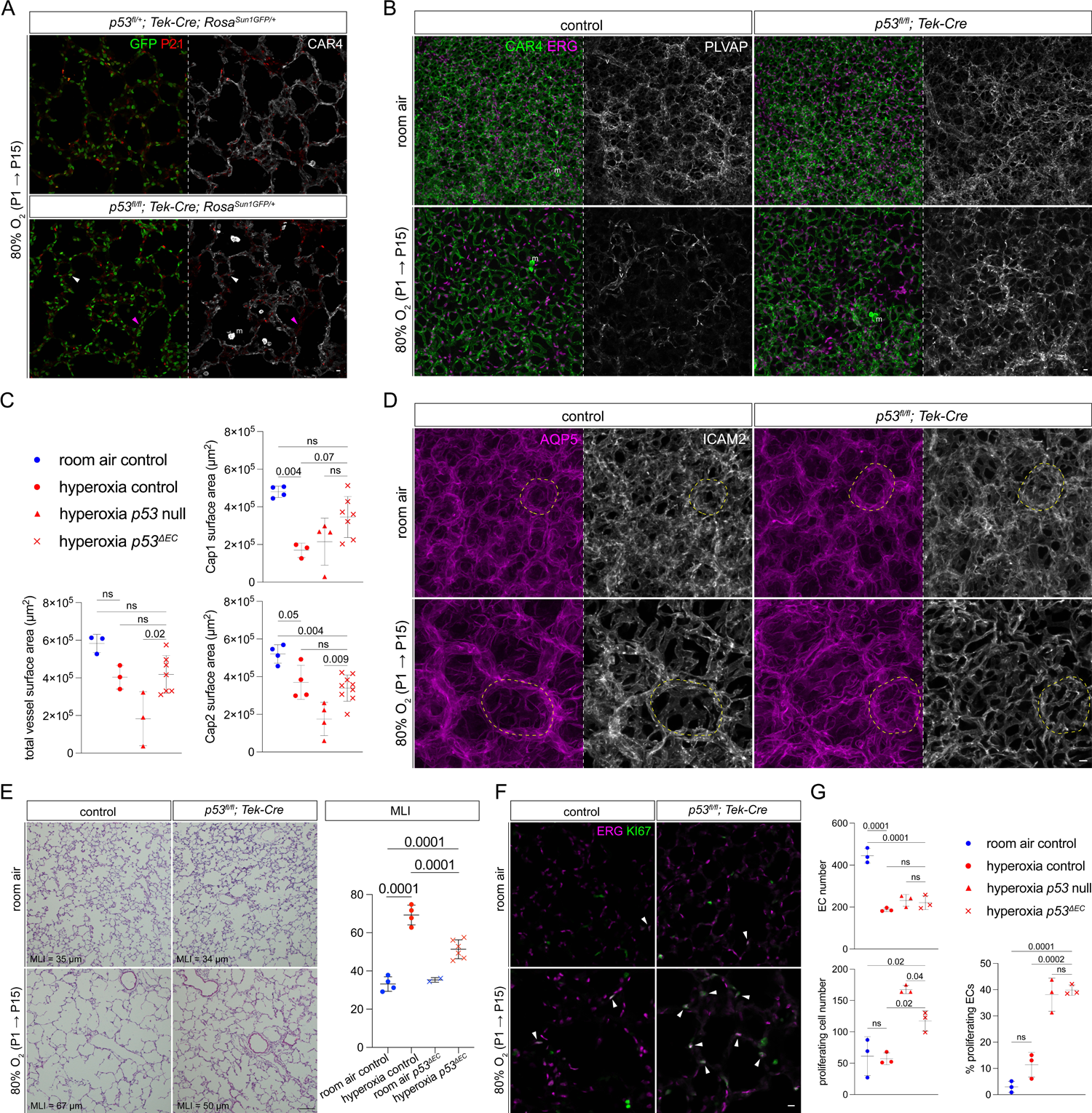
Endothelial-specific *p53* deletion in hyperoxia results in partial rescue of BPD phenotype. (A) Immunostained lung sections showing efficient endothelial deletion of *p53* in ECs using a *Tek-Cre* driver and *Rosa^Sun1GFP^*, as measured by P21 and GFP. Mutant lungs retained occasional P21-expressing escapers in both Cap1 (magenta arrowhead) and Cap2 ECs (white arrowhead). (B) En-face view of immunostained lungs demonstrating the effect of EC-specific *p53* deletion in room air and hyperoxia on EC number (ERG), Cap1 vessels (PLVAP), and Cap2 vessels (CAR4). *p53^ΔEC^* lungs demonstrate a substantial improvement in Cap1 vasculature in hyperoxia compared to control. (**C**) Quantification showing Cap1 vessel area is increased in the hyperoxia *p53^ΔEC^* condition compared to the hyperoxia control, while Cap2 area is relatively unchanged. Compared to the hyperoxia *p53* null, the hyperoxia *p53^ΔEC^* lung shows a significant rescue of Cap2 area and total vessel area (1-way ANOVA with Tukey’s multiple comparisons; ns, not significant). (**D**) En-face view of immunostained lungs demonstrating alveolar islands (circled regions) with improved overall vasculature (ICAM2) in *p53^ΔEC^* hyperoxia, along with a reduction in alveolar simplification (AQP5). (**E**) H&E-stained lung sections from each experimental condition and MLI quantification showing a significant reduction in alveolar simplification in the hyperoxia *p53^ΔEC^* (1-way ANOVA with Tukey’s multiple comparisons). (**F**) Immunostained lung sections showing increased proliferation (KI67) in hyperoxia *p53^ΔEC^* lungs compared to control, mostly in ECs (ERG, white arrowheads). (**G**) Quantification reveals no change in EC number in hyperoxia *p53^ΔEC^* compared to hyperoxia control, and comparable endothelial proliferation between *p53* null and *p53^ΔEC^* lungs (1-way ANOVA with Tukey’s multiple comparisons; ns, not significant). For quantification, each symbol represents the average of 3 distinct regions imaged within 1 mouse lung. P, postnatal. m, macrophage. v, large vessel. Scale bars, 10 um (white bars), 100 um (black bars).

As deletion was confirmed, we continued to analyze the EC-type specific phenotype comparing the *p53^ΔEC^* in room air and hyperoxia (Fig. 6B). Recapitulating the *Cdh5-CreER* model, the vasculature in the *p53^ΔEC^* hyperoxia showed remarkable improvement compared to the hyperoxia control (Fig. 6B). Specifically, Cap1 vessel area had increased and become comparable to Cap1 area at room air, while Cap2 vessel area was maintained compared to hyperoxia control (Fig. 6C). Paradoxically, we detected an increase in Cap2 vasculature and total vessel area comparing *p53* null and *p53^ΔEC^* in hyperoxia (Fig. 6C). Moreover, the alveolar region of the *p53^ΔEC^* mice in hyperoxia showed significant improvement in alveolar angiogenesis and folding of the AT1 cell surface when compared to the control in hyperoxia (Fig. 6D). To quantify changes in airspaces, we measured MLI and found a significant decrease in alveolar size, indicating a partial rescue in the alveolar simplification – a key feature in BPD (Fig. 6E).

To explore whether these changes could be attributed to an upsurge in EC proliferation, we quantified EC number and percentage of proliferating ECs (Fig. 6F). Despite an increase in endothelial proliferation, there was no significant increase in EC number in the *p53^ΔEC^* in hyperoxia when compared to hyperoxia control or *p53* null hyperoxia (Fig. 6G). We then examined DNA damage between hyperoxia control and *p53^ΔEC^*, but found few cells experiencing this process (Fig. S6E). Compared to the hyperoxia *p53* null – which shows an increase in non-EC DNA damage – the endothelial deletion of *p53* in hyperoxia resulted in little γH2AX, indicating that in general, ECs are more resilient to hyperoxic injury than non-ECs.

Given the discrepancy between the *p53* null and *p53^ΔEC^*, particularly (1) the reduction in Cap2 vessel area in the former, (2) the increase in Cap1 vessel area in the latter, (3) the high non-EC expression of γH2AX in the *p53* null, we explored the possibility of non-cell autonomous contributions that could explain these differences. This analysis is particularly relevant since other cell types also upregulate p53 target genes in this context (Fig. S6F). Our previous work highlighted the interplay between the epithelium and the vasculature during development, where Cap2 cells are specified by AT1-specific VEGFA (6). Consequently, we considered the epithelium as a good candidate for this compensation, and deleted *p53* using a pan-epithelial driver, *Shh^Cre^* (28) (Fig. S6G). However, the deletion of *p53* in the epithelium had no effect on the hyperoxia phenotype in the vasculature (Fig. S6G).

### Transitional ECs arise upon conditional deletion of *p53* in the endothelium in hyperoxia

To further explore these conflicting phenotypes, we conducted scRNA-seq of 7447 ECs across the different conditions (Fig. 7A). Since no differences were observed between wild type, *p53* null and *p53^ΔEC^* in room air, only the room air wild type was included in this analysis. For the *p53^ΔEC^* samples in hyperoxia, we sequenced two sets of mice which we refer to as *p53^1EC1^ ^or^ ^2^*, each containing a male and a female. Notably, there was an increase in the early Cap2 cell population, indicating more conversion from Cap1 to Cap2 (Fig. 7A, B, Table S1). Strikingly, a new population of ECs emerged in the *p53^ΔEC^* in hyperoxia consisting of ∼15% of all ECs (Fig. 7A, B). This distinct cell cluster – which we refer to as “transitional EC” – emerged from Cap1 cells and occupies an intermediate transcriptional position between early and mature Cap2 cells, as predicted by Monocle trajectory analysis (29–33) (Fig. 7C). Differential gene analysis between hyperoxia control mice and *p53^ΔEC^* revealed upregulation of some growth factors such as *placental growth factor (Pgf)* and *growth differentiation factor 15 (Gdf15)*, both of which have been implicated in BPD (34–36) (Fig. 7D, Table S4). It also showed upregulation of angiogenesis related genes such as *angiopoetin 2 (Angpt2)*, and genes associated with oxidative stress like *oxidative stress-induced growth inhibitor 1 (Osgin1)* (37, 38) (Fig. 7D).

**Figure 7.**
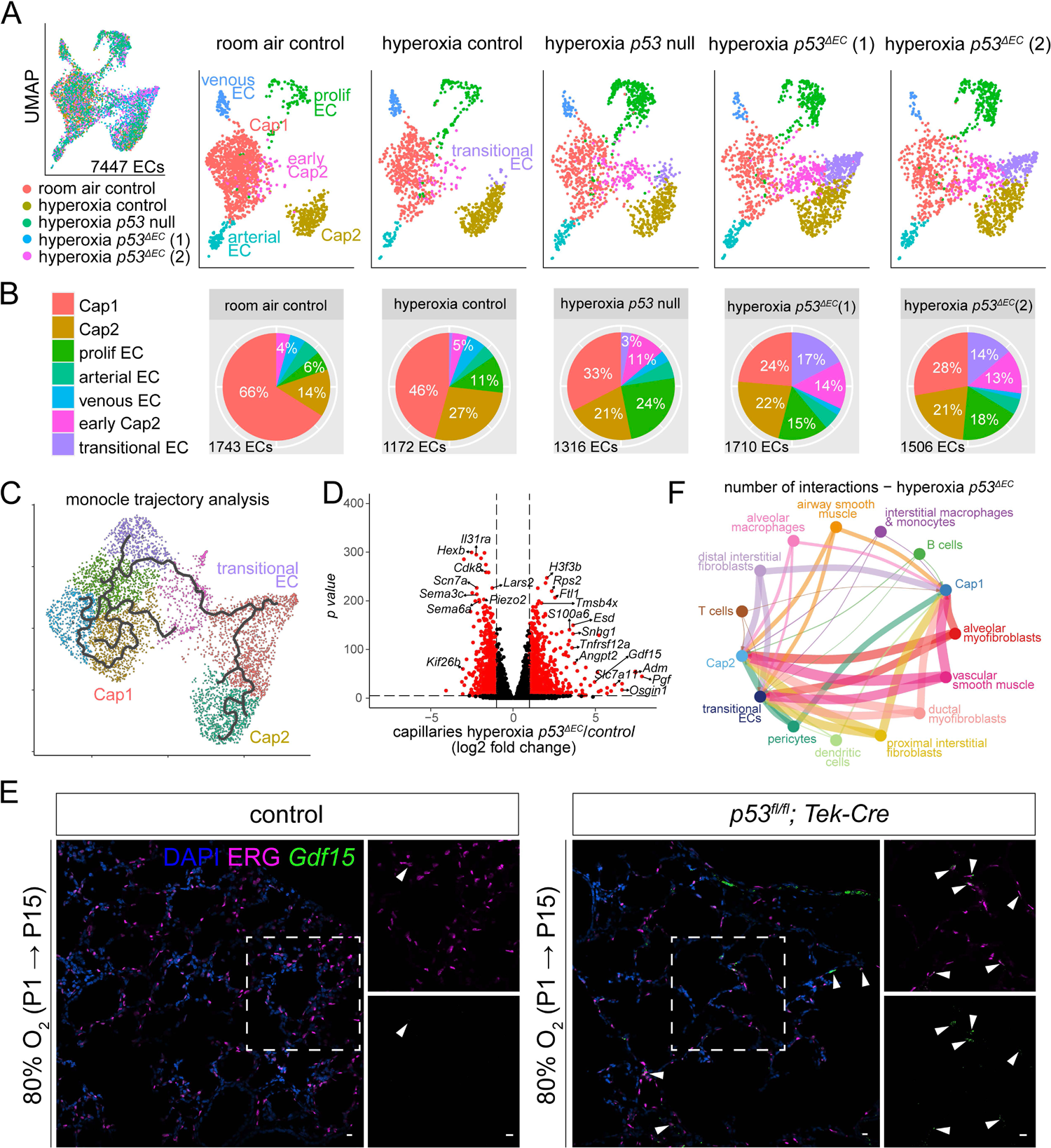
scRNA-seq reveals a novel transitional EC population in the hyperoxia *p53^ΔEC^* lung. (**A**) UMAP of lung ECs, color-coded and overlayed from each experimental condition (left), then split by experimental condition and color-coded according to EC population (right). Notably, the hyperoxia *p53^ΔEC^* lungs demonstrate a novel transitional EC cluster and an increased number of early Cap2 cells. (**B**) Pie charts showing the proportion of individual EC populations in each experimental condition. See Table S1 for complete analysis of cell numbers per population. (**C**) Monocle trajectory analysis UMAP showing the intermediate transcriptional profile of the transitional EC cluster. (**D**) Volcano plot showing differentially expressed genes in the capillaries of the hyperoxia *p53^ΔEC^* lung compared to the hyperoxia control, including the upregulation of *Pgf, Gdf15, Angpt2,* and *Osgin1*. See Table S4 for the complete dataset. (**E**) RNAscope *in situ* hybridization and immunostained lung sections showing upregulation of *Gdf15* in transitional ECs (white arrowheads) in the hyperoxia *p53^ΔEC^* lung compared to the hyperoxia control. (**F**) Circle plot showing ligand-receptor interactions in the hyperoxia *p53^τιEC^* lung with Cap1, Cap2, or transitional ECs as targets. Only mesenchymal and immune interactions with these EC clusters are shown. See Table S5 for the complete dataset. P, postnatal. Scale bars, 10 um.

Gene ontology analysis of the upregulated genes in transitional ECs confirmed these cells might be regulating oxidative and metabolic stress, while regulating other cell populations like fibroblasts (Fig. S7A). Remarkably, despite expressing endothelial lineage markers such as *cadherin-5 (Cdh5) or Platelet endothelial cell adhesion molecule (Pecam1)*, these ECs do not express PLVAP or CAR4, some of the highest Cap1 and Cap2 markers, respectively (Fig. S7B). The early Cap2 cell cluster, which was significantly expanded in the *p53^ΔEC^* mice, does express PLVAP, potentially contributing to the improvement in Cap1 vasculature we observed (Fig. S7B). Additionally, most proliferating cells are Cap1 cells and express PLVAP (Fig. S7B).

To validate the presence of this transitional EC state, we performed RNAscope for *Gdf15* – a specific marker present in the transitional ECs (Fig. S7E), in combination with immunofluorescence for ERG – an EC nuclear marker. Thus, we were able to show significant upregulation of *Gdf15* in ECs in the *p53^ΔEC^* mutant, compared to control hyperoxia (Fig. 7E), consistent with the emergence of a transitional EC. Relevant to these findings, loss of *Gdf15* in neonatal hyperoxia has been shown to result in increased mortality and worsen alveolarization and vascular development, highlighting the importance of transitional EC genes in injury response (35).

We further analyzed these data for *Cdkn1a (p21)* RNA levels in the scRNA-seq data to verify gene deletion and explore the mechanisms for the appearance of this new cluster. Comparison of *p21* across samples revealed that although Cap1 cells in the hyperoxia *p53^ΔEC^* indeed showed similar levels to the Cap1 cells in room air, Cap2 cells and transitional cells had high levels of *p21*, comparable to the control in hyperoxia (Fig. S7C). Additionally, KEGG pathway analysis indicated an upregulation in senescence related genes (Fig. S7D). A possible explanation is that the few escapers we observed (Fig. 6A, S6D) have high expression of *p21,* or that *p21* itself is being activated by an independent signaling mechanism in the *p53^ΔEC^* in hyperoxia. In fact, *p21* has been shown to be regulated by multiple factors outside of *p53*, including *Transforming growth factor β* (*Tgf-*β) (39), growth factors (40), and *activin A* (*Inhba*) (41).

Given that we did not observe changes in the epithelial deletion of *p53*, we considered potential mesenchymal and immune cell contributions, which also upregulate *p53* in hyperoxia (Fig. S6F). To narrow down candidates, we used CellChat in scRNA-seq data to infer cell-cell communication through ligand-receptor interactions (42). Using only Cap1, Cap2 and transitional ECs as targets, we identified interactions arising from mesenchymal and immune cells (Fig. 7F, Table S5). We then compared ligand-receptor interactions between the *p53^ΔEC^* and the *p53* null conditions to detect signals that could explain their conflicting phenotypes (Fig. S8, Table S5). Indeed, we found unique and upregulated interactions in both mouse models upon hyperoxia, with stronger communication between mesenchymal and ECs, and alveolar macrophages on the immune lineage. Interestingly, most of the ligand-receptor interactions were shared for these three EC types; some of these, like *Adrenomedullin - Calcitonin receptor-like receptor (Adm – Calcrl)*, have been associated to EC survival, angiogenesis and inhibition of apoptosis (43). Others, like *Pgf – Vascular endothelial growth factor receptor 1 (Vegfr1), insulin-like growth factor 2 and IGF-1-receptor (Igf2 – Igf1r)* and *Angpt2 – Tek,* have been associated with the regulation of angiogenesis (44, 45). TGF-β, an important pathway in lung development that has been implicated in the pathogenesis of BPD and a *p53*-independent activator of *p21* (39, 46), was one of the shared interactions detected. Overall, this analysis indicates that non-EC contributions could influence vascular survival and repair upon hyperoxia. Furthermore, it suggests that several pro- and anti-angiogenic signals could be modulating Cap1 to Cap2 cell conversion in the *p53^ΔEC^*, as well as Cap1 health, resulting in more Cap1 cells and improved vasculature.

## DISCUSSION

In this study, we demonstrated that neonatal hyperoxia targets Cap2 cells, leading to their demise and subsequent replacement by Cap1 cells, resulting in Cap1 cell depletion and contributing to the vascular rarefaction seen in a mouse model of BPD. Our findings also highlight the pivotal role of the p53 pathway in orchestrating EC regeneration and Cap2 specification during hyperoxic injury-repair. Through global deletion of *p53*, we observed worsened vascular phenotypes, implicating *p53* in mediating vascular recovery. Conversely, endothelial-specific deletion of *p53* reversed the vascular phenotype and improved alveolar simplification, suggesting non-EC contributions to the vascular repair. This recovery was associated with the emergence of a transitional EC state between Cap1 and Cap2 fate, characterized by the upregulation of growth factors and oxidative stress response genes. Our study provides insights into the cellular dynamics underlying vascular injury and repair in BPD, shedding light on the role of EC heterogeneity and the p53 pathway in this context.

Our work contributes to the growing body of literature elucidating the molecular mechanisms underlying BPD pathogenesis. By delineating the cellular dynamics of EC subpopulations and the p53 pathway in vascular injury and repair, we provide novel insights that could inform the development of targeted therapeutic strategies for BPD. Although it has not been tested, it is possible that ablating the p53 pathway in the endothelium could promote vascular resiliency and recovery for both the vasculature and alveolar epithelium in cases of severe BPD.

Moreover, the identification of the transitional EC state expands our understanding of the cellular heterogeneity within the pulmonary vasculature, drawing remarkable comparisons to lung epithelial regeneration and repair. Analogous to the lung epithelium, where damage-associated transient progenitors (DATPs) have been characterized, our study underscores the functional significance of Cap1 cells as endothelial stem-like cells in injury contexts, akin to the role of alveolar type 2 (AT2) cells in epithelial repair (47–49). This parallelism extends to the regulation of cell state by the p53 pathway; DATPs have been described to upregulate *p53* target genes (47–49), and more recently, the role of *p53* in promoting AT1 differentiation has been demonstrated (50), thereby highlighting intriguing similarities in the regulatory mechanisms governing cellular plasticity across different pulmonary cell lineages.

Notably, transitional ECs are not found in wild type or *p53* null lungs exposed to hyperoxia. If transitional ECs do not arise from rare escapers, and are genotypically *p53* mutant cells that activate P21 in a *p53*-independent manner, the question remains as to why we cannot detect them in the *p53* null mice. In this scenario, mesenchymal cell death or stress in the *p53* null might hinder their ability to maintain transitional cells. The underlying ligand-receptor interactions, although predicted by scRNA-seq analysis, need to be elucidated.

While our study provides compelling evidence for the involvement of the p53 pathway in the vascular phenotype of BPD, further studies are warranted to investigate the precise mechanisms by which *p53* regulates EC regeneration and Cap2 specification, including non-cell-autonomous mechanisms. Our data also underlines the interplay between different cell types, and how angiocrine signaling could play a role in injury-repair. The functional significance of the transitional EC state identified in our study requires further investigation to determine its role in vascular repair. Moreover, the conversion of Cap1 to Cap2 fate necessitates a closer look to further dissect the balance between pro- and anti-angiogenic signals that achieve appropriate vascular regeneration.

Additionally, further experiments are needed to better understand the injury mechanism of oxygen and activation of *p53* in this context. Historically, reactive oxygen species (ROS) have been identified as the main culprits for DNA damage in hyperoxia which in turn upregulate *p53* (20, 21, 51). However, changing superoxide levels to decrease oxidative stress has not been sufficient to rescue oxygen toxicity (52). More recent studies emphasize the effects of oxygen in mitochondrial and cellular metabolism, and attribute the damage to degradation of the electron transport chain (ETC) (53). Mitochondrial dysfunction can in turn activate *p53* to reduce induced stress response (ISR) genes; alternatively, ISR genes can themselves induce *p53* (54). It would be interesting to test whether this mechanism is involved in endothelial hyperoxic injury, and if it can be rescued by targeting the different complexes of the ETC.

In conclusion, our findings underscore the importance of EC heterogeneity and *p53* in mediating vascular injury and repair in BPD. By elucidating the cellular dynamics underlying BPD pathogenesis, our study lays the foundation for future research aimed at understanding cell fate choice and regeneration during development and disease.

## METHODS

### Antibodies

The following antibodies were used: rabbit anti-aquaporin5 (AQP5, 1:2500, ab78486, Abcam), mouse carbonic anhydrase IV/CA4 (CAR4, 1:500, AF2414, R&D), rabbit anti-CASPASE3 (CASPASE3, 1:500, 9661, Cell Signaling Technology), PE/Cy7 rat anti-CD45 (1:250, 103114, BioLegend), Alexa Fluor 488 rat anti-CD324 (ECAD, 1:500, 53-3249-80, eBioscience), rabbit anti-avian erythroblastosis virus E-26 (v-ets) oncogene related (ERG, 1:5000, ab92513, Abcam), chicken anti-green fluorescent protein (GFP, 1:5000, AB13970, Abcam), Alexa Fluor 488 mouse anti-H2AX (1:250, BD Biosciences), rat anti-intercellular adhesion molecule 2 (ICAM2, 1:2500, 16-1021-85, eBioscience), Alexa Fluor 647 rat anti-intercellular adhesion molecule 2 (ICAM2, 1:500, A15452, ThermoFisher), rat anti-MKI67 (KI67, 1:1000, 14-5698-82), Cy3-conjugated rat anti-MKI67 (KI67, 1:500, eBioscience, 41-5698-82), rabbit anti-cyclin dependent kinase inhibitor 1A (CDKN1A or P21, 1:1000, 28248-1-AP, ProteinTech), rat anti-plasmalemma vesicle associated protein (PLVAP, 1:125, 553849, BD Biosciences).

### Mouse strains and hyperoxia treatment

All protocols used for this research complied with the Institutional Animal Care and Use Committee regulations of MD Anderson Cancer Center and Northwestern University. The following mouse strains were used (Jackson Lab IDs noted if available): *Trp53^flox/flox^* (24) (008462), *CMV-Cre* (23) (003465), *Tek-Cre* (27) (also called *Tie2-Cre*) (008863), *Cdh5-CreER* (26), *Shh^Cre^* (28) (005622), *Kit^CreER^* (14), *Car4^CreER^* (17) (038602), *Rosa^Sun1GFP^* (15) (021039), and *Rosa^tdT^* (16) (007914). Wild type C57BL/J mice were used to establish a control hyperoxia phenotype and in single-cell RNA sequencing. Mice of both genders were used, and the number of control-mutant pairs are listed in the figure legends. Tamoxifen (T5648; Sigma) dissolved in corn oil (C8267; Sigma) was administered to pups at P0 via intraperitoneal injection to induce Cre-Lox recombination when required; specific doses are noted in the figures. Hyperoxia exposure was performed following published protocols with minor modifications (4). Littermates were either maintained in room air (21% O_2_) or placed in normobaric 80% O_2_ at P0 or P1 if not injected, or P2 if injected, until the date of harvest. A sealed animal chamber (BioSpherix, A30274P) was used to house hyperoxia cages, in which the O_2_ concentration was maintained using an oxygen controller (BioSpherix, P360) connected to an oxygen source. To prevent excessive oxidative stress in the hyperoxia condition, nursing dams were switched between room air and hyperoxia conditions every 24 hours for the duration of exposure.

### Section immunostaining

Lungs were harvested as described in our previous publications (55, 56). Mice were first anesthetized with Avertin (T48402, Sigma), then perfused through the right ventricle with PBS. The trachea was then cannulated, and the lung was inflated using 0.5% paraformaldehyde (PFA, Thermo Scientific, J19943.K2) in PBS at 25 cm H_2_O pressure. After extraction, the lung was fixed in 0.5% PFA at room temperature for 3-6 hours, then briefly washed and left in PBS overnight at 4°C. Section immunostaining was performed as described in our previous publications (55). Fixed lobes were cryoprotected in 20% sucrose in PBS with 10% OCT (Tissue-Tek, 4583) and frozen in OCT blocks. Sections of 20 um thickness were blocked in PBS + 0.3% Triton X-100 and 5% normal donkey serum (Jackson ImmunoResearch, 017-000-121) for one hour. Primary antibodies in PBS + 0.3% Triton X-100 were added to the sections and incubated at 4°C overnight in a humidified container. Sections were washed for 30 minutes in a coplin jar with PBS, then incubated with donkey secondary antibodies and 4’,6-diamidino-2-phenylindole (DAPI) diluted in PBS + Triton X-100 at room temperature for one hour. Following another PBS wash in a coplin jar, sections were mounted with Aqua-Poly/Mount (Polysciences, 18606) and imaged with an Olympus FV1000 confocal microscope using a 30x oil objective.

### Wholemount immunostaining

Wholemount immunostaining was performed as described in our previous publications (6). Thin strips 3 mm in length were cut from the distal edge of the cranial lobe, then blocked in PBS + 0.3% Triton X-100 and 5% normal donkey serum for one hour at room temperature. Strips were added to a solution of primary antibodies diluted in PBS + 0.3% Triton X-100 and rocked overnight at 4°C. The strips were then washed three times in PBS + 0.3% Triton X-100 + 1% Tween-20 (PBSTT) at room temperature for one hour each wash. Secondary antibodies and DAPI were diluted in PBS + 0.3% Triton X-100, and strips were placed in the solution and rocked overnight at 4°C. On the third day, the strips were washed three times in PBSTT as before, then post-fixed in 2% PFA in PBS for two hours. After fixation, the strips were mounted flat side up with Aqua-Poly/Mount and imaged with an Olympus FV1000 confocal microscope. Z-stack images with a step size of 1 um were acquired beginning at the top of the tissue with a depth of 20 um.

### Vasculature analysis and quantification

To analyze and quantify CAR4, PLVAP, and ICAM2 vasculature staining, wholemount immunostained strips were used to ensure comparable, accurate rendering of the vessel surfaces between samples. An Olympus FV1000 confocal microscope was used to image samples with a 30x oil objective and a field size of 424 um x 424 um x 20 um and a pixel dimension of 1024 x 1024 x 20. Three Z-stacks per strip were analyzed with Imaris software to render a surface to measure vascular surface area. To quantity EC number and proliferating cell number, immunostained sections were imaged on the same objective with a field size of 424 um x 424 um x 1 um and a pixel dimension of 1024 x 1024. Imaris surface rendering software was again used to quantify EC number (ERG) and proliferating cell number (KI67) from at least three images per section. Following quantification, GraphPad Prism 10 was used to generate plots and for statistical analysis.

### Mean Linear Intercept (MLI) analysis

MLI was measured on 5 um thick frozen lung sections stained with hematoxylin and eosin (H&E). This quantification was performed in Photoshop in accordance with a published protocol (57). For each mouse lung, three images were acquired on an upright Olympus BX60 microscope with a 10x objective. The guide function of Photoshop was used to add two evenly-spaced vertical and horizontal gridlines on each picture. Along the gridlines, the distance between one alveolar wall and the other was measured using the Photoshop ruler tool. Airways, large vessels, and alveoli in which one wall was not visible were excluded. At least 40 intercepts per image and 3 images per mouse were measured. GraphPad Prism 10 was used to plot MLI values and for subsequent statistical analysis.

### RNAscope *In Situ* hybridization

Harvested lungs were fixed and washed as above and immediately placed in 20% sucrose with 10% OCT (Tissue-Tek, 4583) and 0.5% PFA in PBS overnight at 4°C, then frozen into OCT blocks. Sections of 10 um thickness were probed for *Cdkn1a* and *Gdf15* (Advanced Cell Diagnostics) using the Multiplex Fluorescent Detection Kit v2 (Advanced Cell Diagnostics, 323110). Sections were treated with 4% PFA for 10 minutes, washed with DEPC-H_2_O and 100% ethanol, then treated with hydrogen peroxide for 10 minutes and washed with DEPC-H_2_O. Slides were submerged in boiling 1X target retrieval solution for 4 minutes and treated with Protease III for 30 minutes at room temperature. Probes were added for 2 hours at 40°C and the slides were left in 5x SSC buffer (Thermo Scientific, J60839.K3) overnight. Following a quick wash in RNAscope wash buffer, slides were treated with Amp1 for 30 minutes, Amp2 for 30 minutes, and Amp3 for 15 minutes at 40°C for each treatment with intermittent washes. HRP-C1 was added at 40°C for 15 minutes, then OPAL 520 (1:1500 in TSA buffer, Akoya Biosciences, FP1487001KT) at 40°C for 30 minutes, and finally HRP-blocker for 15 minutes at 40°C with intermittent washes. This same process was repeated for HRP-C2 and OPAL 570 (Akoya Biosciences, FP1488001KT), and for HRP-C3, OPAL 690 (Akoya Biosciences, FP1497001KT). Following RNAscope, sections were immunostained as described above if required. Images were acquired on either an Olympus FV1000 confocal microscope using a 60x oil objective or a Yokogawa CSU-W1 confocal microscope on a Nikon Ti2 Eclipse microscope stand also using a 60x oil objective.

### Cell dissociation and fluorescence-activated cell sorting (FACS)

Lung dissociation was performed as described in our previous publications with minor modifications (6). Whole lungs were harvested in Leibovitz’s Medium (Gibco, 21-083-027), minced with forceps, and digested in Leibovitz’s with 2 mg/mL Collagenase Type I (Worthington, CLS-1, LS004197), 2 mg/mL Elastase (Worthington, ESL, LS002294), and 0.5 mg/mL DNase I (Worthington, D, LS002007) for 30 minutes at 37°C. After 15 minutes of digestion, the tissue was mechanically agitated by pipetting. Fetal bovine serum (FBS, Invitrogen, 10082-139) was then added to a final concentration of 20% and the solution was homogenized. Samples were immediately transferred to the 4°C cold room on ice, where they were filtered with a 70 um cell strainer (Falcon, 352350). Strained samples were centrifuged at 5000 rpm for 1 minute and resuspended in 1 mL of red blood cell lysis buffer (15 mM NH4Cl, 12 mM NaHCO3, 0.1 mM EDTA, pH 8.0) for 3 minutes. Samples were again centrifuged at 5000 rpm for 1 minute, washed with Leibovitz’s + 10% FBS, and filtered into a 5 mL tube with cell strainer cap (Falcon 352235). Cells were then incubated with CD45-PE/Cy7 (BioLegend, 103114), ICAM2-A647 (Invitrogen, A15452), and ECAD-A488 (eBioscience, 53-3249-82) at a concentration of 1:250 for 30 minutes, then centrifuged as above and washed with Leibovitz’s + 10% FBS. Samples were re-filtered and incubated with SYTOX Blue (Invitrogen, S34857), then sorted on a BD FACSAria II Cell Sorter. Excluding dead cells and doublets, four cell populations were collected: CD45+ as the immune lineage, ICAM2+ (from CD45-) as the endothelial lineage, ECAD+ (from CD45- and ICAM2-) as the epithelial lineage, and triple negative representing the mesenchymal lineage.

### Single cell RNA-sequencing

Sequencing and analysis were performed as described and raw data was deposited in GEO (6) (GSEXX pending). FACS-purified cells from each lineage were combined in equal proportions into a single tube and prepared for sequencing using the Chromium Single Cell Gene Expression Solution Platform (10x Genomics). Chromium single-cell RNA-seq output was processed with Cell Ranger and subsequent analysis was performed using R packages Seurat (58) (v5.0.1), Monocle3 (v1.2.9) and CellChat (v1.6.1). Cells were filtered by their gene count to exclude those with counts lower than 200 and higher than 6000, and more than 10% of mitochondrial genes; both parameters were adjusted depending on each sample. We used FindIntegrationAnchors and IntegrateData to aggregate multiple data sets. Cell lineages were established based on the expression of *Cdh1* (epithelium), *Cdh5* (endothelium), *Col3a1* (mesenchyme) and *Ptprc* (immune). Doublets were identified based on the co-expression of these markers and then eliminated. Endothelial cells were subsetted and reclustered and Findmarkers was used to do differential gene analysis between cluster or subpopulations. Enhanced Volcano 1.14.0 was used for plotting with a fold change cutoff of 1 and a p value cut off at 10e-5. Monocle3 and SeuratWrappers were used to analyze capillary ECs and generate pseudotime trajectories. Finally, CellChat objects were created per sample and then merged to perform differential gene expression analysis and extract ligand receptor pairs that were upregulated. For time point comparisons (Fig. S6B), Seurat was used on the published datasets GSE124325 (6).

### Large language models

ChatGPT4 (OpenAI) was used to review and suggest edits for portions of the text, and to troubleshoot coding errors in the R packages used for scRNA-seq analysis.

## ACKNOWLEDGEMENTS

We thank Drs. Guillermina Lozano (The University of Texas MD Anderson Cancer Center, USA), Dieter Saur (Technical University of Munich, Germany) and Ralf Adams (University of Münster, Germany) for providing the *p53^flox^*, *Kit^CreER^* and *Cdh5-CreER* mice, respectively. We thank Celine Kong and Dalia Hassan for assisting with the CellChat analysis and GEO upload. The Advanced Technology Genomics Core and the Flow Cytometry and Cellular Imaging Core Facility were supported in part by The University of Texas MD Anderson Cancer Center and P30CA016672. RNAscope imaging was performed at the Northwestern University Center for Advanced Microscopy supported by NCI CCSG P30CA060553 awarded to the Robert H Lurie Comprehensive Cancer Center. This work was supported by the University of Texas MD Anderson Cancer Center Retention Fund, and National Institutes of Health R01HL130129 and R01HL153511 (JC), and by the Northwestern University Start-up Fund and K99/R00HL155845 (LVE).

## AUTHOR CONTRIBUTIONS

LVE and JC designed the research; LVE and JB performed the research; LVE and JB wrote the paper; all authors read and approved the paper.

