## Supplemental Figures for "Endothelial deletion of *p53* generates transitional endothelial cells and improves lung development during neonatal hyperoxia"

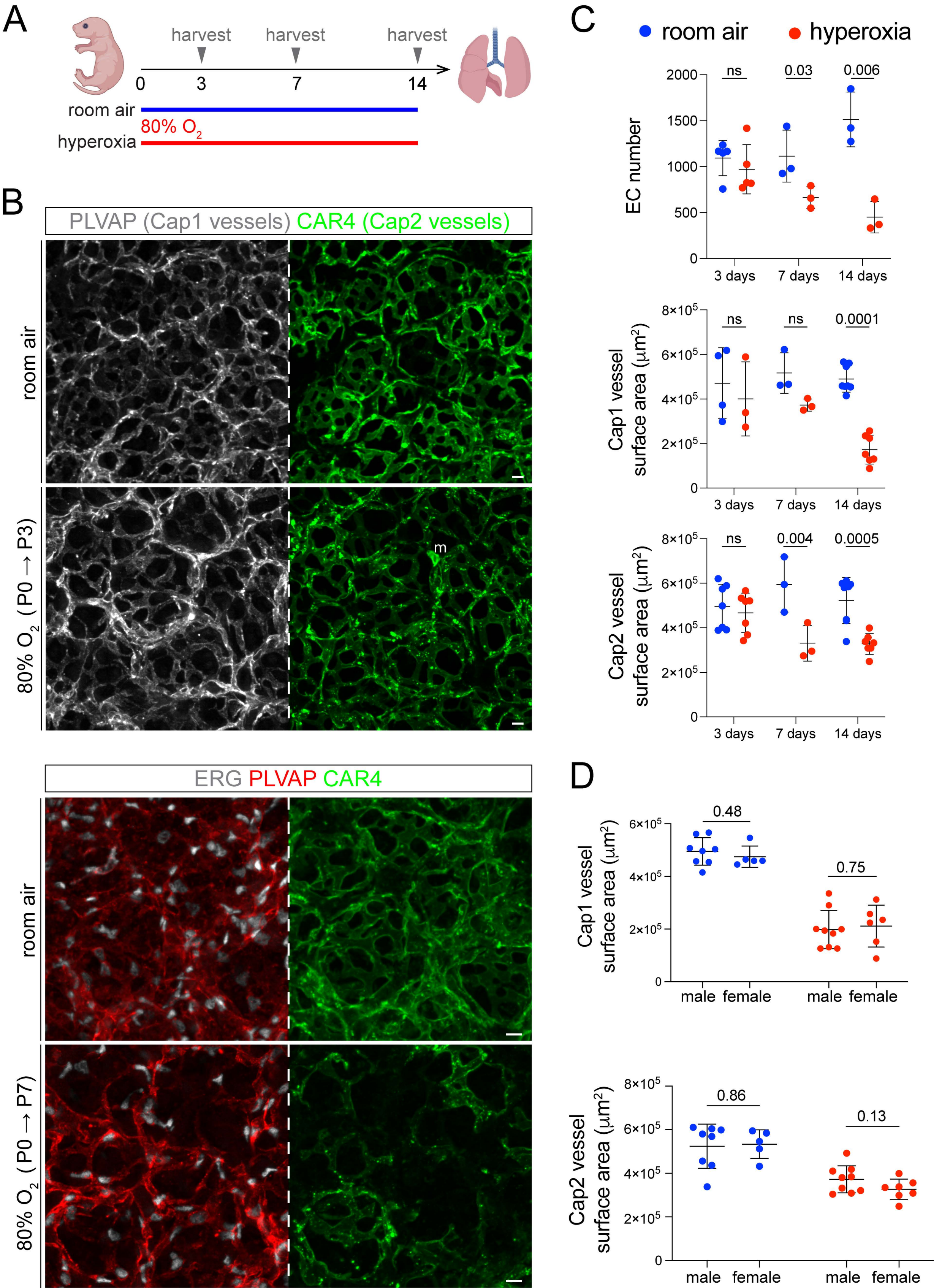
A
harvest
harvest
harvest
0
3
7
14
room air
80% O2
hyperoxia
B
PLVAP (Cap1 vessels) CAR4 (Cap2 vessels)
room air
80% O2 ( P0 → P3)
m
ERG PLVAP CAR4
room air
80% O2 ( P0 → P7)
C
room air
hyperoxia
EC number
ns
0.03
0.006
3 days
7 days
14 days
Cap1 vessel surface area (μm2)
ns
ns
0.0001
Cap2 vessel surface area (μm2)
ns
0.004
0.0005
D
Cap1 vessel surface area (μm2)
0.48
0.75
male
female
male
female
Cap2 vessel surface area (μm2)
0.86
0.13
male
female
male
female

**Figure S1**

(**A**) Experimental model showing continuous neonatal exposure to hyperoxia (80% $O_2$) or room air for 3, 7, or 14 days. Created with BioRender.com. (**B**) En-face view of immunostained lungs showing the effect of 3 days and 7 days of hyperoxia exposure on EC number (ERG), Cap1 vasculature (PLVAP), and Cap2 vasculature (CAR4) compared to room air control. (**C**) Quantification of EC number, Cap1 area, and Cap2 area after 3, 7, or 14 days of hyperoxia exposure and room air controls. 3 days of hyperoxia results in no significant changes, 7 days of exposure results in significantly reduced EC number and Cap2 area with no significant change in Cap1 area, while 14 days of exposure results in a significant reduction in EC number, Cap1 area, and Cap2 area (Student's t test; ns, not significant). (**D**) Quantification of Cap1 area and Cap2 area in room air and hyperoxia after 14 days of exposure divided into male and female mice. No significant sex-dependent differences in vessel area were observed in any condition (Student's t test). Images are representative of at least 3 littermate pairs. For quantification, each symbol represents the average of 3 distinct regions imaged within 1 mouse lung. P, postnatal. m, macrophage. Scale bars, 10 um.

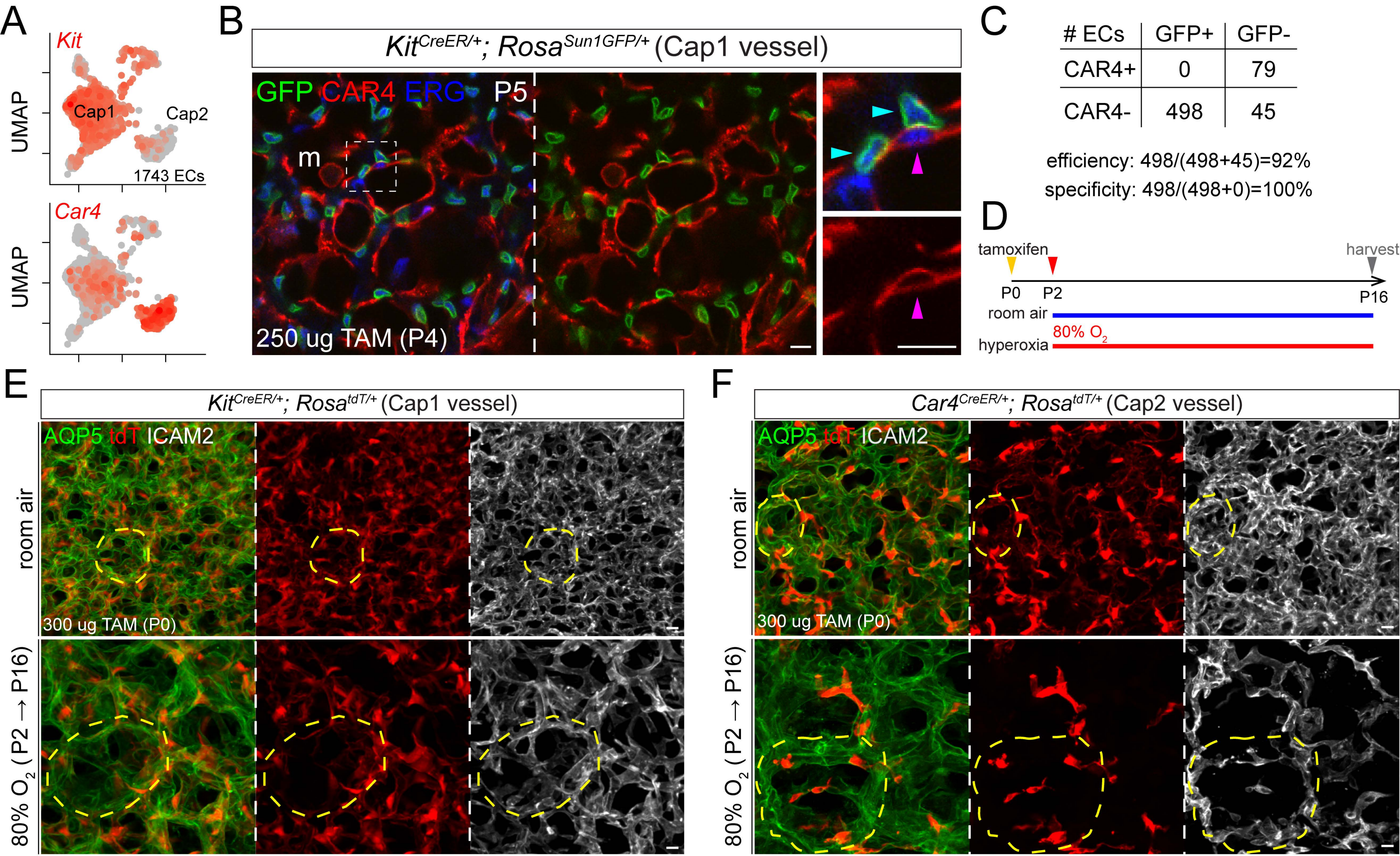
A
Kit
Cap1
Cap2
1743 ECs
UMAP
Car4
UMAP
B
KitCreER/+; RosaSun1GFP/+ (Cap1 vessel)
GFP CAR4 ERG P5
m
250 ug TAM (P4)
C
### ECs | GFP+ | GFP-
CAR4+ | 0 | 79
CAR4- | 498 | 45
efficiency: 498/(498+45)=92%
specificity: 498/(498+0)=100%
D
tamoxifen
harvest
P0
P2
P16
room air
80% O2
hyperoxia
E
KitCreER/+; RosatdT/+ (Cap1 vessel)
AQP5 tdT ICAM2
room air
300 ug TAM (P0)
80% O2 (P2 → P16)
F
Car4CreER/+; RosatdT/+ (Cap2 vessel)
AQP5 tdT ICAM2
room air
300 ug TAM (P0)
80% O2 (P2 → P16)

**Figure S2**

(**A**) UMAP of purified lung ECs showing expression of *Kit* and *Car4* as markers for Cap1 and Cap2 ECs, respectively. (**B**) Immunostained lung sections showing the efficiency and specificity of the $Kit^{CreER}$ driver. Cap1 ECs are labeled with GFP, while Cap2 ECs (CAR4 positive) lack GFP expression. (**C**) Quantification of the efficiency and specificity of the $Kit^{CreER}$ driver. Approximately 92% of all Cap1 ECs were labeled using the driver, while 100% of GFP labeled cells were Cap1 ECs. (**D**) Experimental model showing tamoxifen administration at P0 in EC type-specific drivers to label Cap1 ECs ($Kit^{CreER}$) or Cap2 ECs ($Car4^{CreER}$) with fluorescent protein tdTomato (tdT) for lineage tracing, followed by 14 days of hyperoxia exposure beginning at P2. Created with BioRender.com. (**E-F**) En face view of immunostained lungs showing the alveolar (AQP5) and vascular (ICAM2) simplification in hyperoxia, confirming that tamoxifen injection combined with the $Kit^{CreER}$ driver (**E**) or $Car4^{CreER}$ driver (**F**) does not impact the phenotype. Images are representative of at least 3 littermate pairs. P, postnatal. m, macrophage. Scale bars, 10 um.

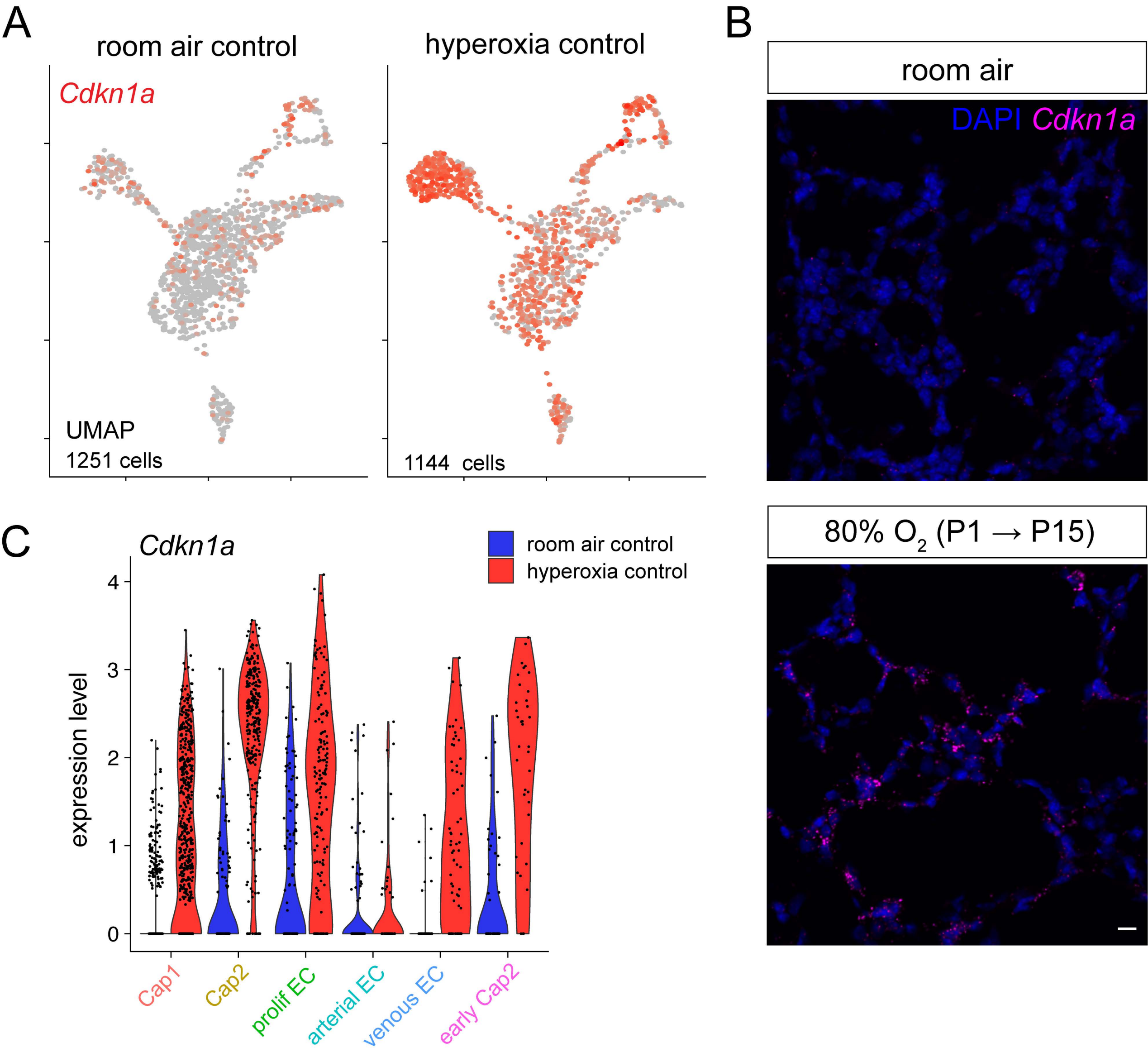
A
room air control
hyperoxia control
Cdkn1a
UMAP
1251 cells
1144 cells
B
room air
DAPI Cdkn1a
80% O2 (P1 → P15)
C
Cdkn1a
room air control
hyperoxia control
expression level
0
1
2
3
4
Cap1
Cap2
prolif EC
arterial EC
venous EC
early Cap2

**Figure S3**

(**A**) UMAP of purified lung capillary ECs showing expression of *p53* target gene *Cdkn1a* in room air and hyperoxia, demonstrating the upregulation of *Cdkn1a* in lung ECs in hyperoxia. (**B**) RNAscope *in situ* hybridization of lung sections showing the widespread upregulation of *Cdkn1a* in hyperoxia compared to room air. (**C**) Violin plot showing expression of *Cdkn1a* in each EC population in room air and hyperoxia. *Cdkn1a* is upregulated in every EC population except arteries. Images are representative of at least 3 littermate pairs. P, postnatal. Scale bars, 10 um.

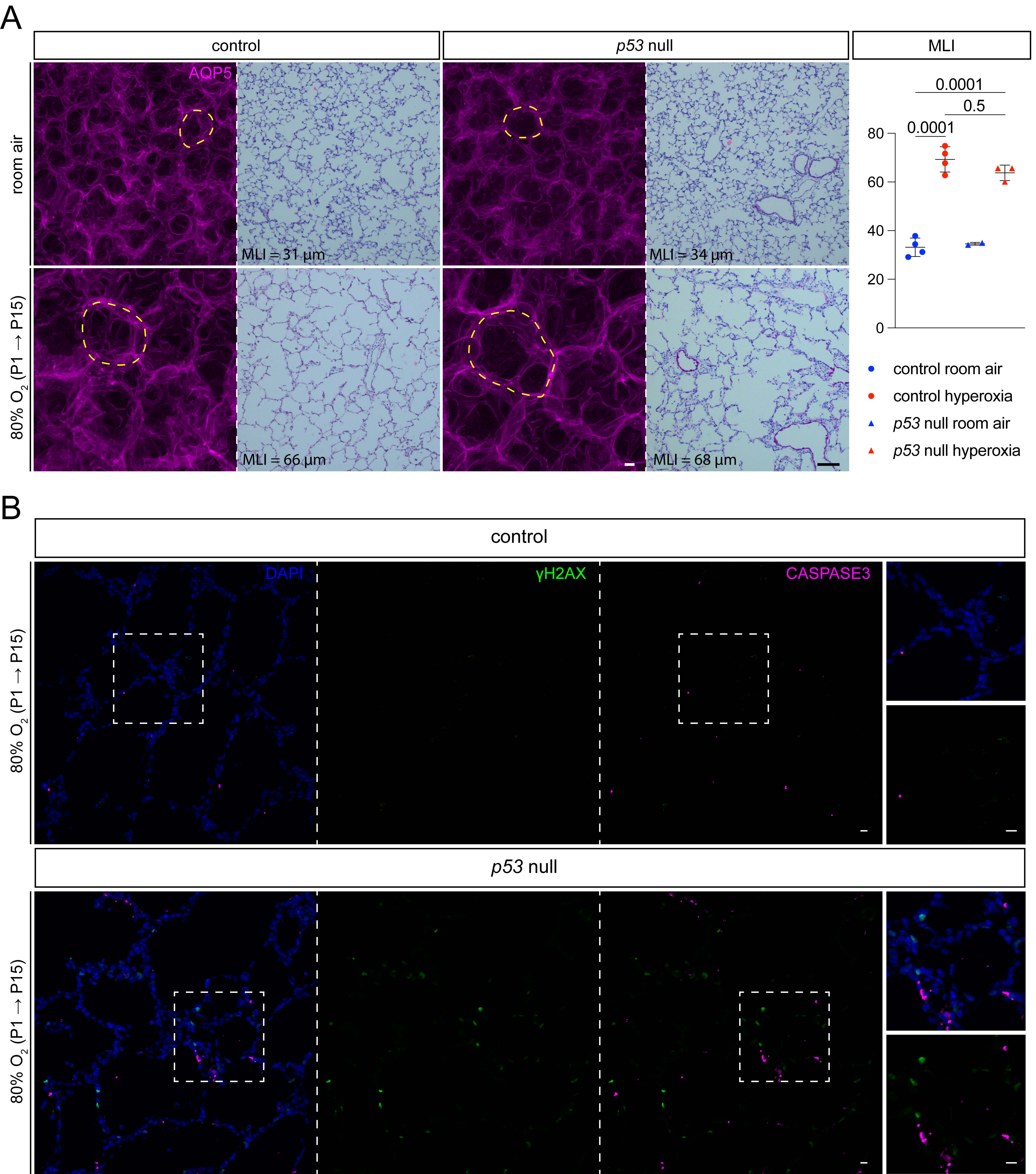
A
control
p53 null
MLI
AQP5
room air
MLI = 31 µm
MLI = 34 µm
80% $O_2$ (P1 → P15)
MLI = 66 µm
MLI = 68 µm
0.0001
0.5
0.0001
80
60
40
20
0
control room air
control hyperoxia
p53 null room air
p53 null hyperoxia
B
control
DAPI
γH2AX
CASPASE3
80% $O_2$ (P1 → P15)
p53 null
80% $O_2$ (P1 → P15)

**Figure S4**

(**A**) En-face view of immunostained lungs and H&E-stained lung sections showing the extent of alveolar simplification in each experimental condition, along with associated MLI quantification. AQP5 staining reveals a similar degree of simplification between hyperoxia control and hyperoxia *p53* null, with comparable enlargement of alveolar islands (dashed regions). MLI quantification confirms airspace enlargement is comparable between hyperoxia control and hyperoxia *p53* null (Student's t test). Each symbol represents the average of 3 distinct regions imaged within 1 mouse lung. (**B**) Immunostained lung sections showing expression of γH2AX and cleaved CASPASE3 in hyperoxia control and *p53* null lungs. γH2AX and CASPASE3 are both upregulated in the *p53* null compared to control, but little co-localization was observed between markers. Boxed regions are magnified. Images are representative of at least 3 littermate pairs. P, postnatal. Scale bars, 10 um (white bars), 100 um (black bars).

A

Gene list for Cap2 cell score

| | |
|---|---|
| Itgb5 | Trim16 |
| Chst1 | Tmem47 |
| Sirpa | Tbx3 |
| Tbx2 | Agfg1 |
| Car4 | Ly6a |
| Ednrb | Prx |
| Emp2 | Ly6c1 |
| Apln | Timp3 |
| Rtn1 | Mgst3 |
| Enho | Stxbp6 |
| Pmp22 | Arhgef3 |
| Tppp3 | Tspan15 |
| Igfbp7 | Rgs12 |
| Cd24a | Rcsd1 |
| Tbxa2r | Bcam |
| Kitl | Pdlim2 |
| Cyp4b1 | Tmcc2 |
| Gem | Mgll |
| Rasgrp2 | Atp13a2 |
| Nhlrc2 | Dhrs3 |
| Pvrl3 | Nrp1 |
| Kdr | Lhfp |
| Plaur | Mpped2 |
| Ptp4a3 | Pcdh1 |

B

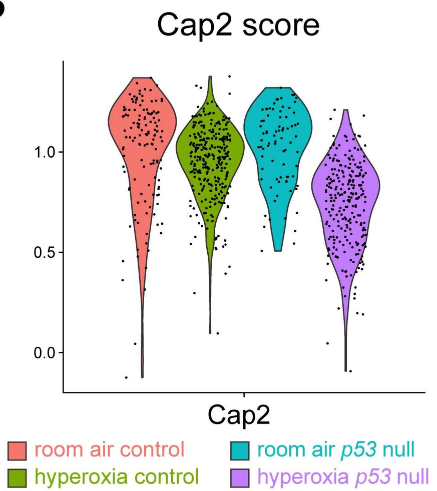


C

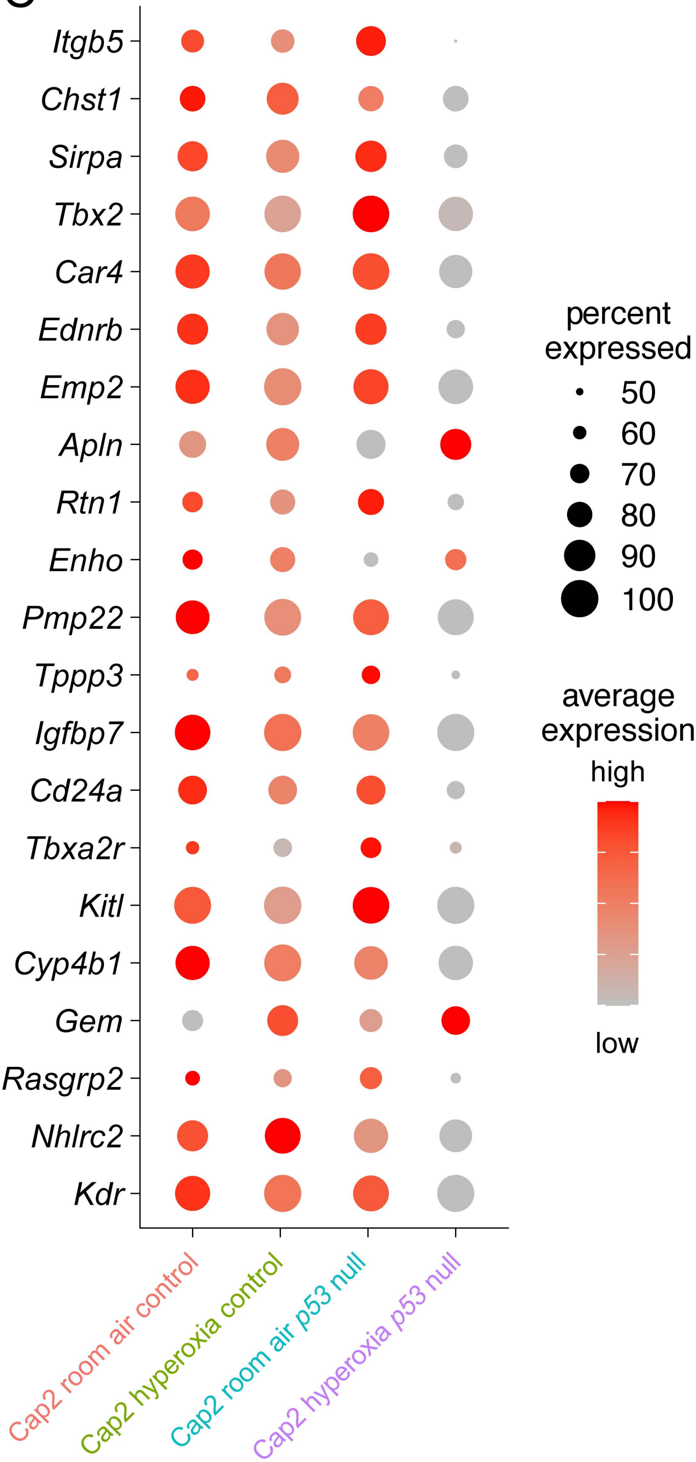

**Figure S5**

(**A**) Gene list of Cap2 EC marker genes expressed at room air at P14 as determined by scRNA-seq that were used to build a Cap2 score. (**B**) Violin plot showing the Cap2 score for each condition which suggests Cap2 cells significantly downregulate their marker genes in the hyperoxia *p53* null compared to all other conditions. (**C**) Dot plot showing the expression of the top 22 Cap2 genes in each condition which are substantially downregulated in the *p53* null lung in hyperoxia.

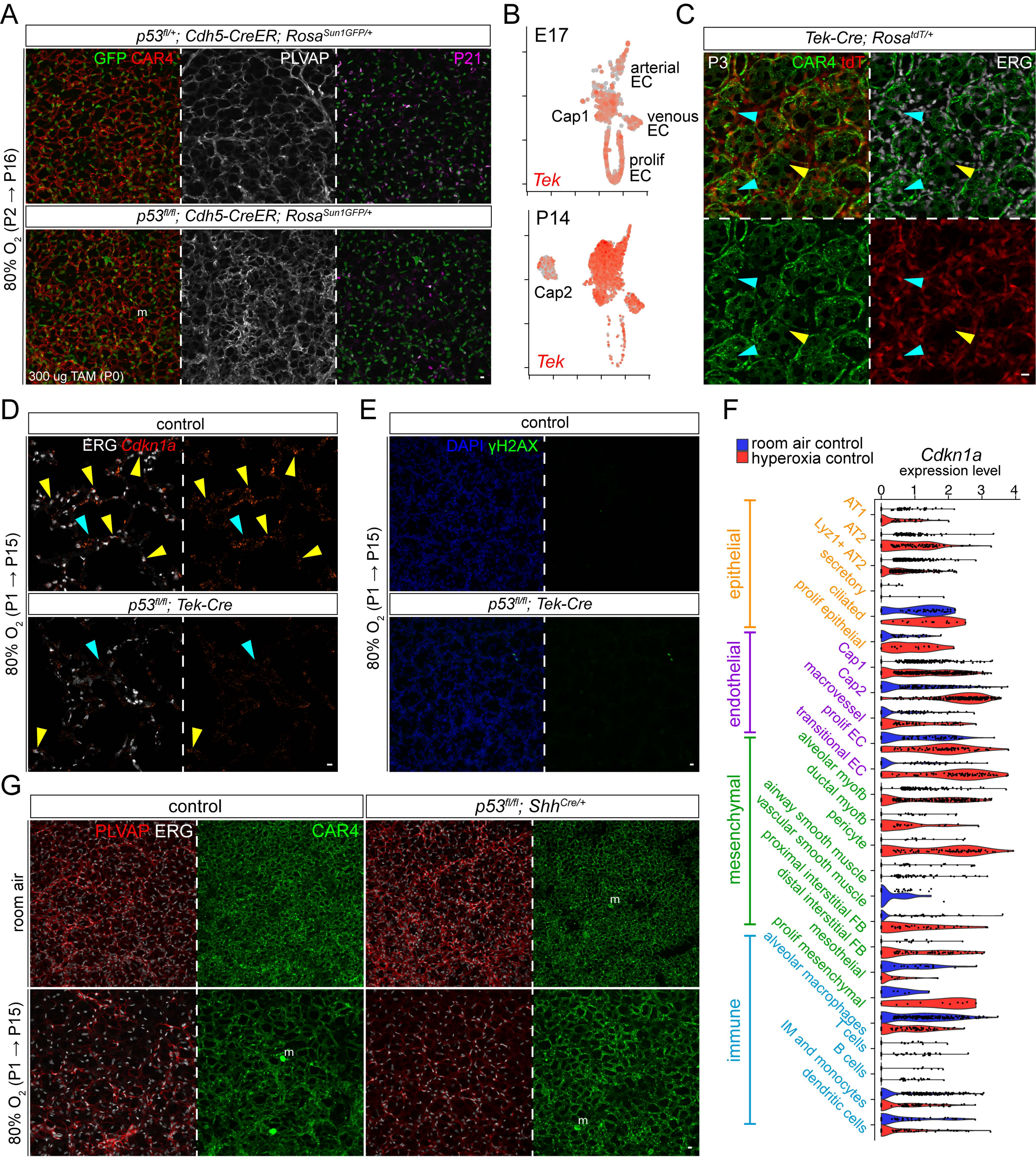

A
p53fl/+; Cdh5-CreER; RosaSun1GFP/+
GFP CAR4
PLVAP
P21
p53fl/fl; Cdh5-CreER; RosaSun1GFP/+
80% O2 (P2 → P16)
300 ug TAM (P0)
m
B
E17
arterial EC
Cap1
venous EC
prolif EC
Tek
P14
Cap2
Tek
C
Tek-Cre; RosatdT/+
P3
CAR4 tdT
ERG
D
control
ERG Cdkn1a
p53fl/fl; Tek-Cre
80% O2 (P1 → P15)
E
control
DAPI γH2AX
p53fl/fl; Tek-Cre
80% O2 (P1 → P15)
F
room air control
hyperoxia control
Cdkn1a
expression level
0 1 2 3 4
epithelial
endothelial
mesenchymal
immune
AT1
AT2
Lyz1+ AT2
secretory
ciliated
prolif epithelial
Cap1
Cap2
macrovessel
prolif EC
transitional EC
alveolar myofb
ductal myofb
pericyte
airway smooth muscle
vascular smooth muscle
proximal interstitial FB
distal interstitial FB
mesothelial
prolif mesenchymal
alveolar macrophages
T cells
B cells
IM and monocytes
dendritic cells
G
control
p53fl/fl; ShhCre/+
PLVAP ERG
CAR4
room air
80% O2 (P1 → P15)
m

**Figure S6**

(**A**) En-face view of immunostained lungs showing the EC type-specific effect of *p53* deletion using the pan-endothelial *Cdh5-CreER* driver in combination with the *Rosa*$^{Sun1GFP}$ reporter. Endothelial deletion of *p53* resulted in an overall improvement in vascular density compared to the control in hyperoxia, most notably in the Cap1 vasculature (PLVAP). This mutant showed incomplete deletion of P21 as assessed by GFP+ and P21+ cells. (**B**) UMAP of purified lung ECs showing expression of *Tek* at E17 and P14. Though the postnatal Cap2 population expresses low levels of *Tek*, it is expressed at E17 in the immature Cap1 population (or progenitors) which gives rise to the Cap2 cells. (**C**) En-face view of immunostained lungs showing expression of tdT in lineage-traced ECs using the *Tek-Cre* driver. ERG staining revealed the driver to be highly specific to ECs, with most Cap2 ECs showing expression of tdT (cyan arrowheads) and rare Cap2 escapers lacking tdT expression (yellow arrowhead). (**D**) RNAscope *in situ* hybridization and immunostained lung sections showing the expression levels of *Cdkn1a* in the hyperoxia control and the hyperoxia *p53*$^{\Delta EC}$ conditions. The *p53*$^{\Delta EC}$ lung exhibited a significant reduction in *Cdkn1a*-expressing ECs (ERG, yellow arrowheads), but non-EC *Cdkn1a* expression (cyan arrowheads) was conserved. (**E**) En-face view of immunostained lungs showing expression of γH2AX in the hyperoxia control and hyperoxia *p53*$^{\Delta EC}$ lungs, showing similar levels of DNA damage in the hyperoxia *p53*$^{\Delta EC}$ lung compared to control. (**F**) Violin plot showing expression levels of *p53* target gene *Cdkn1a* across cell populations from each cell lineage in the lung, showing widespread upregulation of *Cdkn1a* in hyperoxia in nearly every population. (**G**) En-face view of immunostained lungs showing the deletion of *p53* in the epithelium using the *Shh*$^{Cre}$ driver, which demonstrates no vascular phenotype in room air or hyperoxia, as assessed by CAR4, PLVAP, and ERG staining. Images are representative of at least 3 littermate pairs. P, postnatal. AT1, alveolar type 1. AT2, alveolar type 2. EC, endothelial cell. myofb, myofibroblast. FB, fibroblast. IM, interstitial macrophage. m, macrophage. Scale bars, 10 um.

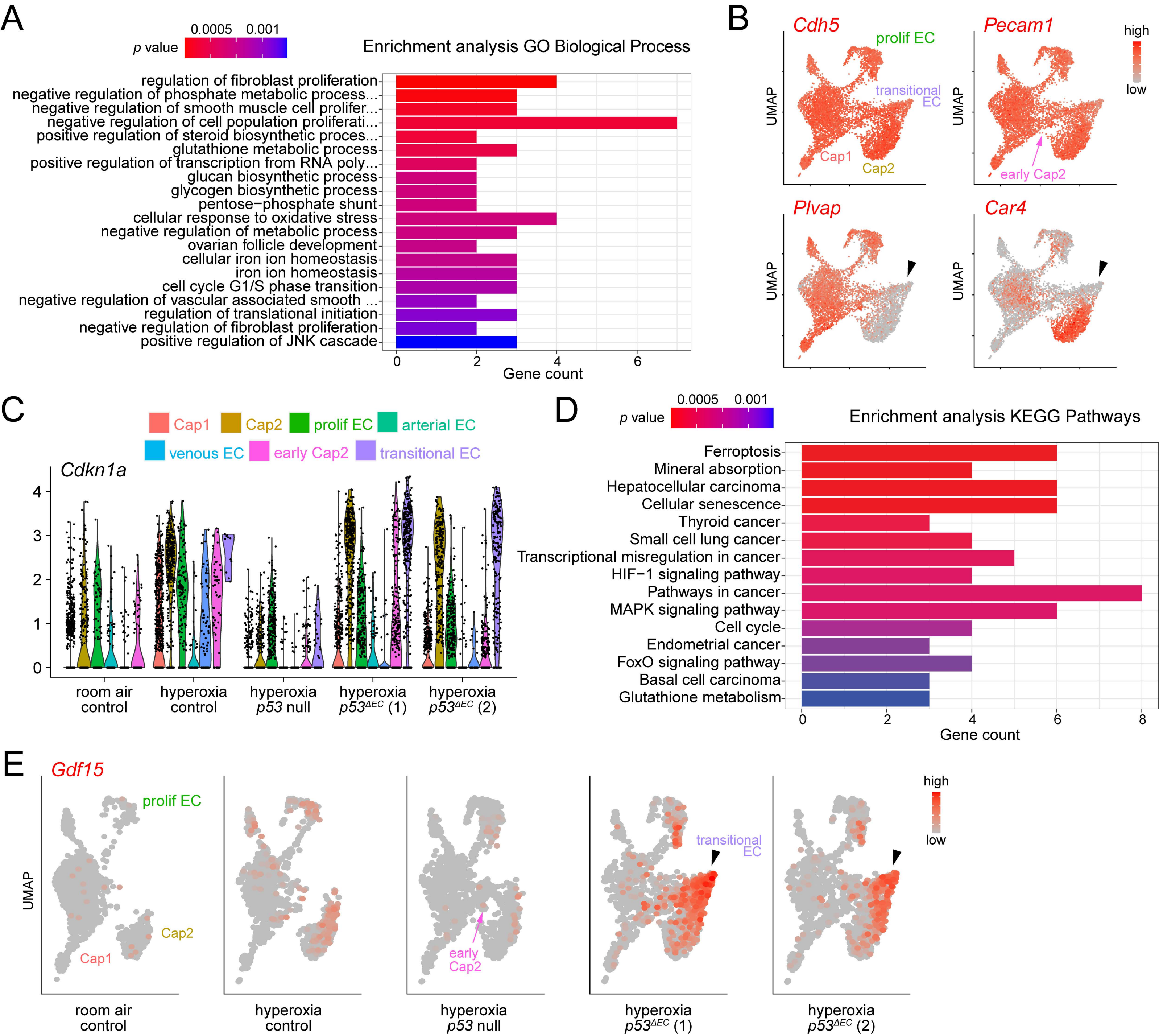

A
p value
0.0005 0.001
Enrichment analysis GO Biological Process
regulation of fibroblast proliferation
negative regulation of phosphate metabolic process...
negative regulation of smooth muscle cell prolifer...
negative regulation of cell population proliferati...
positive regulation of steroid biosynthetic proces...
glutathione metabolic process
positive regulation of transcription from RNA poly...
glucan biosynthetic process
glycogen biosynthetic process
pentose−phosphate shunt
cellular response to oxidative stress
negative regulation of metabolic process
ovarian follicle development
cellular iron ion homeostasis
iron ion homeostasis
cell cycle G1/S phase transition
negative regulation of vascular associated smooth ...
regulation of translational initiation
negative regulation of fibroblast proliferation
positive regulation of JNK cascade
Gene count
B
Cdh5
Pecam1
Plvap
Car4
prolif EC
transitional EC
Cap1
Cap2
early Cap2
UMAP
high
low
C
Cap1
Cap2
prolif EC
arterial EC
venous EC
early Cap2
transitional EC
Cdkn1a
room air control
hyperoxia control
hyperoxia p53 null
hyperoxia p53ΔEC (1)
hyperoxia p53ΔEC (2)
D
p value
0.0005 0.001
Enrichment analysis KEGG Pathways
Ferroptosis
Mineral absorption
Hepatocellular carcinoma
Cellular senescence
Thyroid cancer
Small cell lung cancer
Transcriptional misregulation in cancer
HIF−1 signaling pathway
Pathways in cancer
MAPK signaling pathway
Cell cycle
Endometrial cancer
FoxO signaling pathway
Basal cell carcinoma
Glutathione metabolism
Gene count
E
Gdf15
prolif EC
Cap1
Cap2
early Cap2
transitional EC
UMAP
high
low
room air control
hyperoxia control
hyperoxia p53 null
hyperoxia p53ΔEC (1)
hyperoxia p53ΔEC (2)

**Figure S7**

(**A**) Gene ontology analysis showing the upregulated biological processes in transitional ECs indicating a role in the response to oxidative and metabolic stress, along with the regulation of other cell populations. (**B**) UMAPs of purified hyperoxia *p53*$^{\Delta EC}$ lung ECs showing expression of EC markers *Cdh5* and *Pecam1*, and EC-type specific markers *Plvap* and *Car4*. Notably, transitional ECs (arrowhead) express both lineage endothelial markers, but lack expression of *Plvap* and *Car4*. Early Cap2 cells express *Plvap* but not *Car4*. (**C**) Violin plots showing expression levels of *Cdkn1a* across EC populations in each condition. *p53*$^{\Delta EC}$ lungs in hyperoxia exhibit substantial expression of *Cdkn1a* in both Cap2 ECs and transitional ECs at RNA level compared to hyperoxia control. (**D**) KEGG pathway enrichment analysis showing an upregulation of senescence related genes in transitional ECs. (**E**) UMAP of lung ECs showing the expression of *Gdf15* across conditions, and its specific upregulation in transitional ECs (arrowhead).

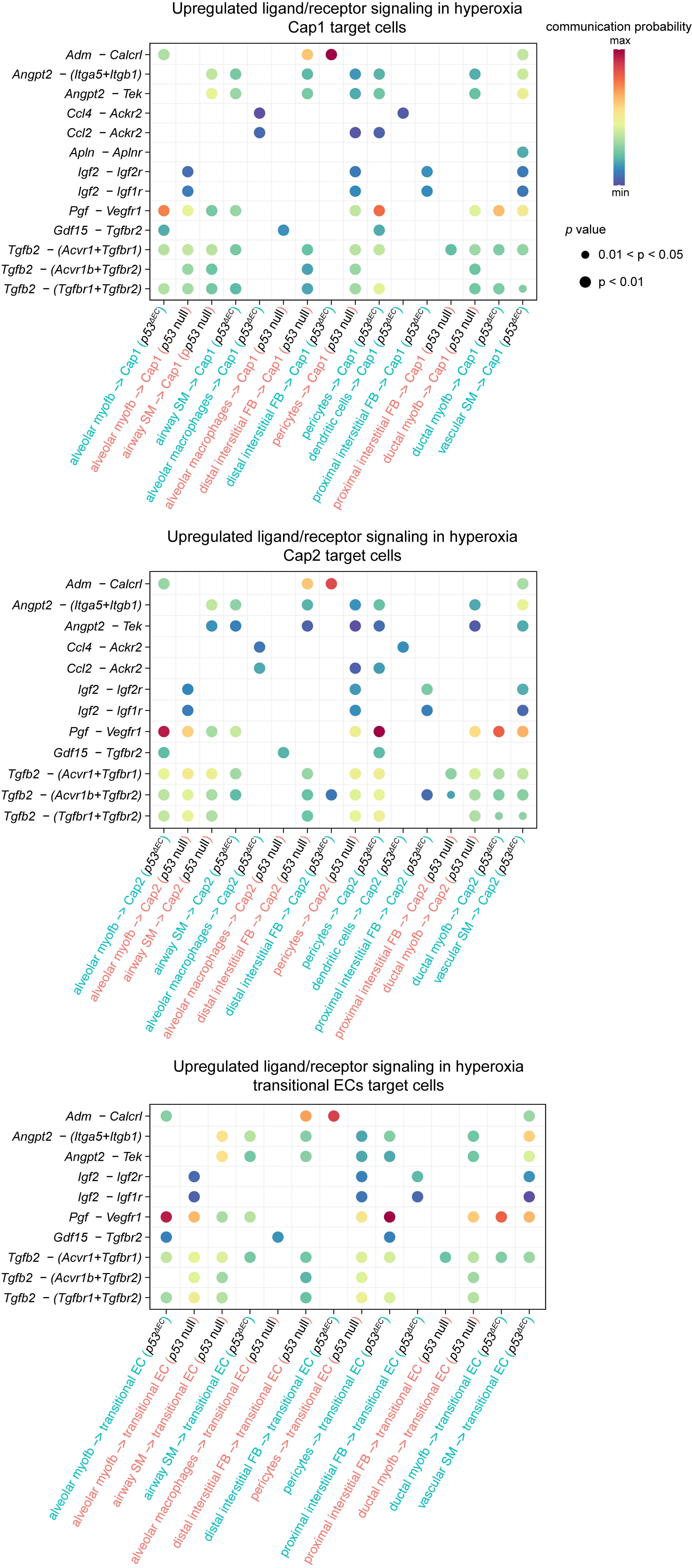
Upregulated ligand/receptor signaling in hyperoxia
Cap1 target cells
Adm – Calcrl
Angpt2 – (Itga5+Itgb1)
Angpt2 – Tek
Ccl4 – Ackr2
Ccl2 – Ackr2
Apln – Aplnr
Igf2 – Igf2r
Igf2 – Igf1r
Pgf – Vegfr1
Gdf15 – Tgfbr2
Tgfb2 – (Acvr1+Tgfbr1)
Tgfb2 – (Acvr1b+Tgfbr2)
Tgfb2 – (Tgfbr1+Tgfbr2)
alveolar myofb –> Cap1 (p53ΔAEC)
alveolar myofb –> Cap1 (p53 null)
airway SM –> Cap1 (p53 null)
airway SM –> Cap1 (p53ΔAEC)
alveolar macrophages –> Cap1 (p53ΔAEC)
alveolar macrophages –> Cap1 (p53 null)
distal interstitial FB –> Cap1 (p53 null)
distal interstitial FB –> Cap1 (p53ΔAEC)
pericytes –> Cap1 (p53 null)
pericytes –> Cap1 (p53ΔAEC)
dendritic cells –> Cap1 (p53ΔAEC)
proximal interstitial FB –> Cap1 (p53ΔAEC)
proximal interstitial FB –> Cap1 (p53 null)
ductal myofb –> Cap1 (p53 null)
ductal myofb –> Cap1 (p53ΔAEC)
vascular SM –> Cap1 (p53ΔAEC)
communication probability
max
min
p value
0.01 < p < 0.05
p < 0.01
Upregulated ligand/receptor signaling in hyperoxia
Cap2 target cells
Adm – Calcrl
Angpt2 – (Itga5+Itgb1)
Angpt2 – Tek
Ccl4 – Ackr2
Ccl2 – Ackr2
Igf2 – Igf2r
Igf2 – Igf1r
Pgf – Vegfr1
Gdf15 – Tgfbr2
Tgfb2 – (Acvr1+Tgfbr1)
Tgfb2 – (Acvr1b+Tgfbr2)
Tgfb2 – (Tgfbr1+Tgfbr2)
alveolar myofb –> Cap2 (p53ΔAEC)
alveolar myofb –> Cap2 (p53 null)
airway SM –> Cap2 (p53 null)
airway SM –> Cap2 (p53ΔAEC)
alveolar macrophages –> Cap2 (p53ΔAEC)
alveolar macrophages –> Cap2 (p53 null)
distal interstitial FB –> Cap2 (p53 null)
distal interstitial FB –> Cap2 (p53ΔAEC)
pericytes –> Cap2 (p53 null)
pericytes –> Cap2 (p53ΔAEC)
dendritic cells –> Cap2 (p53ΔAEC)
proximal interstitial FB –> Cap2 (p53ΔAEC)
proximal interstitial FB –> Cap2 (p53 null)
ductal myofb –> Cap2 (p53 null)
ductal myofb –> Cap2 (p53ΔAEC)
vascular SM –> Cap2 (p53ΔAEC)
Upregulated ligand/receptor signaling in hyperoxia
transitional ECs target cells
Adm – Calcrl
Angpt2 – (Itga5+Itgb1)
Angpt2 – Tek
Igf2 – Igf2r
Igf2 – Igf1r
Pgf – Vegfr1
Gdf15 – Tgfbr2
Tgfb2 – (Acvr1+Tgfbr1)
Tgfb2 – (Acvr1b+Tgfbr2)
Tgfb2 – (Tgfbr1+Tgfbr2)
alveolar myofb –> transitional EC (p53ΔAEC)
alveolar myofb –> transitional EC (p53 null)
airway SM –> transitional EC (p53 null)
airway SM –> transitional EC (p53ΔAEC)
alveolar macrophages –> transitional EC (p53 null)
distal interstitial FB –> transitional EC (p53 null)
distal interstitial FB –> transitional EC (p53ΔAEC)
pericytes –> transitional EC (p53 null)
pericytes –> transitional EC (p53ΔAEC)
proximal interstitial FB –> transitional EC (p53ΔAEC)
proximal interstitial FB –> transitional EC (p53 null)
ductal myofb –> transitional EC (p53 null)
ductal myofb –> transitional EC (p53ΔAEC)
vascular SM –> transitional EC (p53ΔAEC)

**Figure S8**

DotPlot of predicted ligand-receptor interactions that are upregulated in the *p53*$^{\Delta EC}$ lung ECs. Using Cap1 (top), Cap2 (middle) and transitional ECs (bottom) as targets, we compared potential communications between mesenchymal and immune populations in the *p53*$^{\Delta EC}$ or *p53* null. Although most of the interactions were shared between ECs, differences between mutants are present which could possibly explain the endothelial differences observed in both models. See Table S5 for the complete dataset. Myofb, myofibroblast. EC, endothelial cell. SM, smooth muscle. FB, fibroblast.
